# RNA polymerase II CTD Ser5 phosphorylation induces competing effects of expansion and compaction

**DOI:** 10.1101/2025.09.11.675598

**Authors:** Michaela R. Cohen, Wei Chen, Sophia M. Dewing, William P. Barr, Kian A. Sethi, Ray B. East, Oluebube C. Onwuzulu, Scott A. Showalter, K. Aurelia Ball

**Affiliations:** Skidmore College, Department of Chemistry, 815 N Broadway, Saratoga Springs, NY 12866, United States; Center for Eukaryotic Gene Regulation, Department of Biochemistry and Molecular Biology, The Pennsylvania State University, 107 Althouse Laboratory, University Park, PA 16802, United States; Department of Chemistry, The Pennsylvania State University, 104 Benkovic Building, 376 Science Drive, University Park, PA 16802, United States; Aldevron, 4055 41st Avenue South Fargo, North Dakota 58104, United States; Department of Chemistry, Johns Hopkins University, 138 Remsen Hall, 3400 N. Charles Street, Baltimore, MD 21218, United States; Department of Biochemistry, University of Wisconsin, Madison, 413 Bock Laboratories, 1525 Linden Drive, Madison, WI 53706, United States; Department of Chemistry & Biochemistry, University of California, Merced, 5200 North Lake Rd., Merced, CA 95343, United States

## Abstract

The carboxy-terminal domain (CTD) of RNA Polymerase II, composed of tandem heptad repeats with the consensus sequence YSPTSPS, orchestrates the transcription cycle through a dynamic series of post-translational modifications. Among these, the phosphorylation of Ser5 is critical for initiator/promoter clearance and the recruitment of capping enzymes. However, the exact conformational consequences of these modifications are still not fully understood. This study investigates how Ser5 phosphorylation affects the local and global conformation of the CTD, its influence on proline isomerization, and how variations in the repeat sequence modulate these effects. We employed Gaussian accelerated Molecular Dynamics (GaMD) simulations on 3-heptad models of both the consensus CTD sequence and an Asn7 variant. We found that Ser5 phosphorylation promotes expansion of the peptide due to the repulsion between the negatively-charged phosphate groups, but also increases the population of *cis*-Pro6, which leads to compaction. We used a clustering algorithm to identify commonly populated conformations, with a focus on those conformations that change in population with Ser5 phosphorylation. Our simulations reveal that the expansion of the CTD due to Ser5 phosphorylation is accompanied by a change in local, intra-heptad interactions in both variants. Notably, phosphorylation significantly increases the population of *cis*-Pro6 due to steric repulsion between the Asn7 side chain and the large side chain of the phosSer5, but has a smaller increase in the consensus variant. These results clarify the underlying mechanisms by which phosphorylation can modulate the CTD’s structural landscape to regulate the transcription cycle.

**Significance:** The RNA Polymerase II CTD is a critical part of the machinery that regulates transcription, and therefore, understanding how it functions in this process is essential. However, the conformational effects of known modifications to the CTD, such phosphorylation and proline isomerization, are not fully understood. This paper uses all-atom molecular dynamics simulations to identify the specific conformational changes to the disordered CTD with phosphorylation, and with changing heptad sequence. We also identify the interactions that are responsible for these changes. Our results emphasize that two chemical properties of phosphate groups, their negative charge and their large size, can affect protein conformation. For the CTD, these properties have competing effects on the overall compaction of the disordered sequence.

## Introduction

The RNA Polymerase II (RNAP II) Carboxy-Terminal Domain (CTD) is involved in coordinating various steps of the transcription cycle, including initiation, promoter proximal pausing, elongation and splicing (1). The intrinsically disordered CTD consists of a repeat of a heptad of residues (Fig. 1), the number of which varies across different species. Post-translational modifications at different residues in the CTD heptad sequence help regulate the key steps in the transcription process through what is known as the ‘CTD code’. As the heptad repeats contain two conserved proline residues, proline *cis*-*trans* isomerization is also believed to represent a component of the CTD code that can influence transcription (1). Currently, it is believed that the CTD is hypo-phosphorylated when assembling the initiation complex but then needs to be phosphorylated at Ser5 to release RNAPII into elongation mode by dissociating from the Mediator complex (1). Given the disordered nature of the CTD, and the large number of nearly identical heptad repeats, it has been challenging to understand how phosphorylation affects the CTD at the structural level, mediating its functional role. Early clostripain cleavage, gel filtration, and sucrose gradient ultracentrifugation studies suggested that the CTD is compact when unphosphorylated and more extended when phosphorylated (2, 3). However, more recent small-angle X-ray scattering (SAXS) experiments indicated no difference in the overall radius of gyration with phosphorylation for an 11-repeat segment of the *Drosophila melanogaster* CTD (4, 5). Additionally, NMR spectroscopy revealed that phosphorylation at Ser5 increased the population of *cis* conformation at Pro6, especially in variants of the heptad sequence that contained Asn rather than Ser at position seven (4). This shows that sequence differences in the CTD heptad repeats could help regulate transcription by influencing the proline residue conformation, but currently the structural details of how variations in sequence and phosphorylation state affect proline conformation are not well understood.

**Figure 1.**
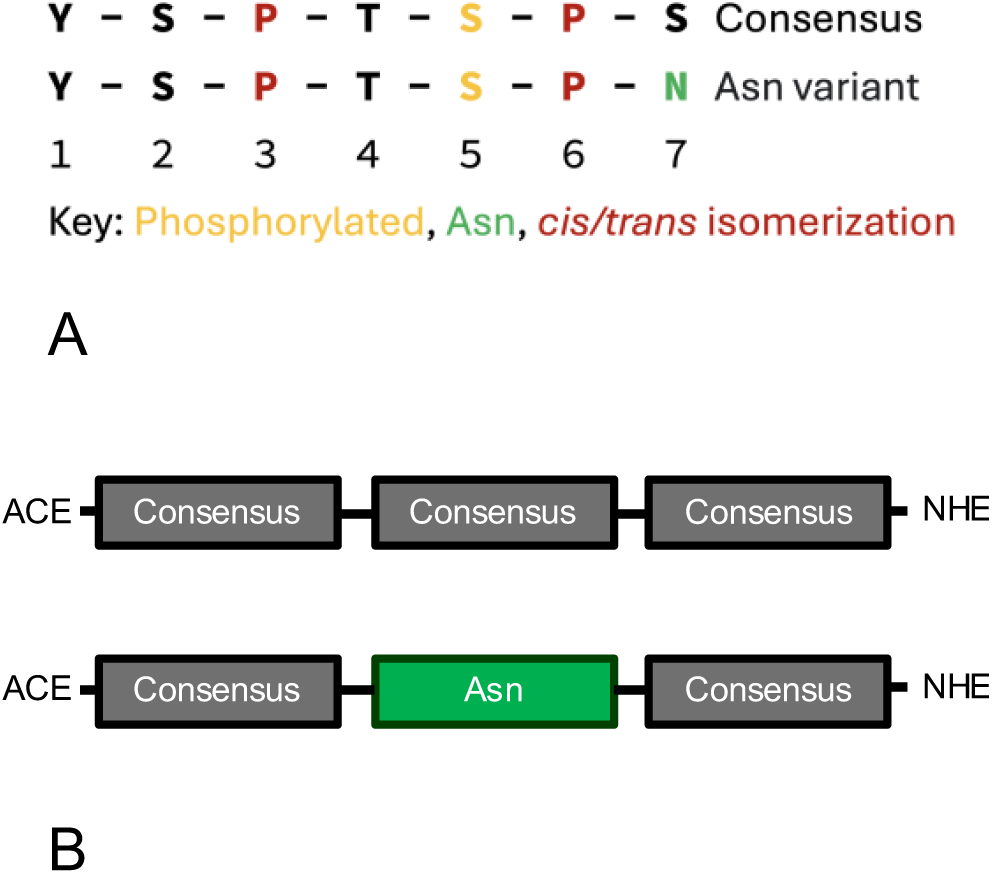
(*A*) Heptad sequences used in our simulations and experiments. (B) Peptide sequences consisted of 3 repeats of the heptad sequence with protecting groups capping the N and C termini. The first and last heptads were the consensus in all constructs, while the middle heptad was either the consensus or Asn variant. In phosphorylated simulations, Ser5 in all three heptads was phosphorylated.

**Figure 2.**
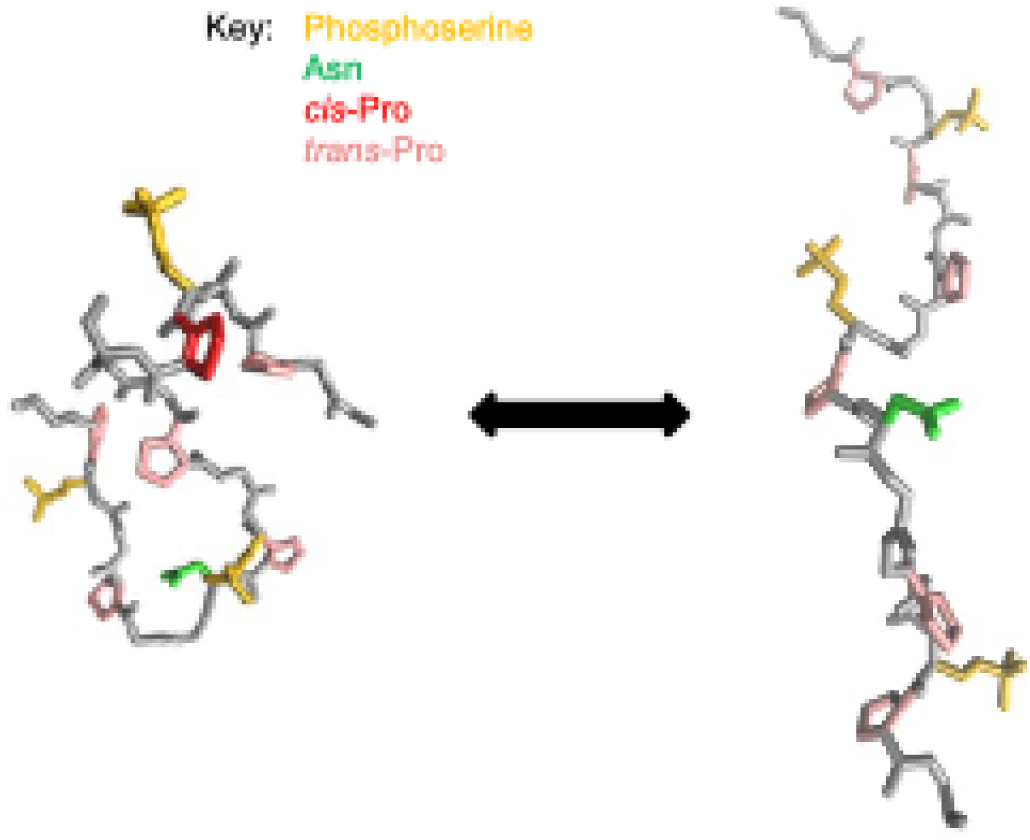
Visualization of global conformational change in CTD structure. Asn residues are shown in green, phosphoserine in yellow, *cis*-Pro in red, and *trans*-Pro in pink.

For most amino acids, only the *trans* conformation of the peptide bond is energetically accessible due to steric repulsion between the α carbons in the *cis* conformation; however, proline can access both *cis* and *trans* conformations due to its unique ring structure that includes the backbone nitrogen. In folded proteins in the Protein Data Bank, 5-7% of prolines are stabilized in the *cis* state (6). Intrinsically disordered proteins (IDPs) occupy an ensemble of conformations, and these sequences are often rich in proline residues, which occupy an equilibrium between the *cis* and *trans* conformations, where the *cis* population is typically in the range of 5-20% of the ensemble (7–10). Given the ability of disordered sequences to switch between *cis* and *trans* populations on a slow timescale (seconds to minutes) compared to cellular processes, switching the proline conformational state can serve as a mechanism for cellular regulation (1, 11, 12). Both the termination factor Nrd1 and the phosphatase Ssu72 are known to preferentially bind the CTD phosSer5 sequence with Pro6 in a *cis* conformation (1). Proline isomerases, like Ess1, can therefore influence the binding process by facilitating the transition between the *cis* and *trans* states (1, 12). The conformation of the proline peptide bond is also context dependent, allowing proline to play different roles in different IDPs. Previous work has shown that the identity of neighboring residues can affect the *cis* population for prolines, both in the context of folded protein structures and in short, disordered segments (13, 14). While high proline content is typically associated with a more expanded IDP ensemble (15), the *cis* conformation creates a kink in the peptide backbone, introducing more compact conformations into the overall conformational ensemble of a proline-rich IDP. Simulations have shown that *cis*-Pro conformations disrupt secondary structure and lead to more compact ensembles (16, 17). As with the sequence of neighboring residues, the phosphorylation state of adjacent residues can affect proline’s tendency to adopt a *cis* conformation (10, 18). For the CTD, phosphorylation could allow the *cis* conformation and the associated conformational changes to be modulated dynamically in the cell.

The effect of phosphorylation on the IDP conformational ensemble has been previously studied as a potential mechanism of regulation, as many IDPs contain phosphorylation sites. Phosphorylation affects the chemistry of a serine residue by adding a –2 charge and by increasing the volume of the residue by 55% (19). Due to the increase in negative charge, hyperphosphorylation has generally been observed to expand neutral and negatively-charged IDPs and collapse positively-charged IDPs (19, 20). However, as with the CTD, there are other IDPs that show little change in average radius of gyration with phosphorylation. For Sic1 and Ash1, phosphorylation changes the local structure in different ways, increasing some types of interactions and decreasing others, but maintains the overall chain expansion (21–23). In the case of Ash1, the mix of charged residues and prolines in the sequence is believed to be responsible for a lack of change in overall compaction with phosphorylation (22). In some cases, the steric effects of the large, hydrophilic side chain can also lead to expansion by disrupting clustering of hydrophobic residues (24), while in other cases phosphorylation can lead to increased hydrophobic interactions (23, 25). The structural effects of phosphorylation detected by NMR are primarily local to the residues near the phosphorylation site (22, 26). As in the case of the CTD, these effects can include shifting the equilibrium of *cis* and *trans* conformations for nearby proline residues. In the case of p53, phosphorylation of Ser46 eliminates the *cis*-Pro47 isomer (10). Martin et al. found that for Ash1, phosphorylation does not affect the overall *cis* proline populations in the protein, but there are small changes for particular prolines near some phosphorylation sites (22). Although many IDPs are both proline-rich and regulated by phosphorylation, little is currently understood about the specific mechanisms by which the addition of a phosphate group can affect the conformation of nearby prolines and change the *cis* propensity.

Here, we employ all-atom molecular dynamics (MD) simulations to investigate how Ser5 phosphorylation affects the local and global conformation of the CTD in atomistic detail. In order to make the simulations more tractable, we focus on a short peptide consisting of a three-heptad consensus repeat (Fig. 1). Due to the repeating nature of the CTD sequence, local effects of phosphorylation should be evident even in such a short peptide. The middle heptad is in a similar sequence environment to a heptad that is part of the longer CTD, and will be able to form interactions with its neighboring heptads. Variants of the heptad sequence with an asparagine in position 7 showed larger effects of phosphorylation on proline conformation in previous NMR studies of the 11-heptad CTD, motivating us to perform additional simulations on a construct with Asn at position 7 only in the middle heptad (Asn variant) (Fig. 1). To overcome the large energy barrier between the *cis*-Pro and *trans*-Pro conformations and sample the transition in our simulations, we employed Gaussian accelerated MD (GaMD) simulations. We complement our computational results with SAXS experiments performed on the same peptide constructs used in the simulations. Our simulations are consistent with experimental data and show that phosphorylation has competing effects on the CTD conformational ensemble. While the repulsion between negatively-charged phosSer5 residues drives a slight expansion in the ensemble, steric clashes between the phosphate group and neighboring side chains drives an increase in the *cis*-Pro6 conformation, counteracting this expansion. This effect is enhanced for the Asn variant due to the larger Asn side chain.

## Methods

### MD Simulations

MD simulations were run on a Linux cluster using Amber 18 pmemd CUDA for GPU functionality on four constructs: S5S7, pS5S7, S5N7, pS5N7, as summarized in Table 1 (27). GaMD was used to overcome the barrier between the *cis* and *trans* states of the *⍵* angle for enhanced sampling (28). Each construct consisted of a 3-heptad repeat of the consensus sequence, except for the S5N7 and pS5N7 constructs, which substituted Asn7 in place of Ser7 for only the middle heptad. S5S7 and S5N7 are both unphosphorylated constructs, whereas pS5S7 and pS5N7 are phosphorylated at Ser5 of each heptad. For each construct, two different starting structures for GaMD simulations were generated with Monte Carlo (MC) simulations using the CAMPARI software package (29). MC simulations were performed using the OPLS-AA/L force field in a spherical droplet with a 100 Å radius with the ABSINTH implicit solvation model. Sodium chloride was modeled explicitly and set at 10 mM. Replica exchange was performed with 20 replicas at temperatures spanning from 230 to 420 K for 51,000,000 steps. The first 1,000,000 steps were discarded as equilibration time.

**Table 1.** Summary of simulations run.

| Construct | # of simulations | Length analyzed per simulation<br>(ns) | Total length<br>analyzed (μs) |
| --- | --- | --- | --- |
| S5S7 | 10 | 480 | 4.8 |
| pS5S7 | 10 | 480 | 4.8 |
| S5N7 | 10 | 490 | 4.9 |
| pS5N7 | 10 | 490 | 4.9 |

Each independent simulation was run for between 500-600 ns, but only the last 480 ns were analyzed for the consensus variant, and the last 490 ns for the Asn variant. When running the simulations, we used the Amber 18 software package with the CUDA implementation of PMEMD. We used the Amber 18 LeAP module to create topology and coordinate files. All simulations used the *TIP3P-FB* water model for solvation and the *ff99SB* force field in a truncated octahedral box, with the box’s borders placed at least 15 Å from all atoms in the peptide. To best address the electrostatic effects of the phosphate groups, we used previously published force field parameters (30, 31). To neutralize charges due to the phosphate groups, 6 Na+ ions were added. We performed two sets of minimization steps, first restraining the peptide with a harmonic force potential of 10 kcal/mol/Å^2^, and the other set without restraints. The system was then heated from 100 to 300 K with a timestep of 2 fs for a total of 40 ps using a Langevin thermostat (γ = 1.0), with a 10 kcal/mol/Å^2^ restraint on the peptide. Next, three stages of equilibration were performed, first with a Berendsen barostat using a 10 kcal/mol/Å^2^ restraint on the peptide, with pressure and density converging to 1.013 bar for a total of 50 ps. A second unrestrained round of equilibration was then performed for 500 ps using a Monte Carlo barostat. Finally, we performed a third round of equilibration with parameters specific to GaMD, as in a previous study (16), for 52 ns. A final production was run for 490-600 ns under isobaric conditions with a 2 fs time step and a 0.1 ps frame output for reweighting. The last 480-490 ns of each simulation was used for analysis. Bonds to hydrogen were constrained using the SHAKE algorithm. The particle-mesh Ewald procedure was used to handle long-range electrostatic interactions with a non-bonded cutoff of 9 Å for the direct space sum.

### Trajectory Analysis

All simulations were analyzed using a combination of the *cpptraj* module in AmberTools 24 (32) and in-house Python scripts for data processing and plotting. Structural figures were created using Visual Molecular Dynamics (VMD) software (33).

The simulated ensemble was reweighted to correct for the GaMD free energy boost. We determined appropriate weights for each frame in the trajectories by comparing potential of mean force (PMF) plots for the proline *ω* dihedral angle using both the cumulant expansion on the second order and Maclaurin series expansion to the 10th order provided by the PyReweighting tool kit, as in a previous study (16). The Maclaurin expansion method was determined to be most useful as it allowed data reweighting independent of the reaction coordinate.

The *cis* and *trans* conformations were defined as in a previous study: *cis* (*ω* = –90° to +50°) and *trans* (*ω* = +100° to +240°) (16).

*P*(*r*) plots were created by measuring pair-wise distances between α carbons for every 100 frames in the simulation trajectories (every 10 ps). Both simulation and experimental *P*(*r*) plots were scaled by setting the maximum value to 1.

### Error Calculations

The error bars in Figs. 3, 5, 7, 8, 9 *E-F* were calculated using a formula for a weighted standard error of the mean. The error was calculated by first determining the effective sample size, *N*:

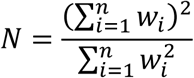

**Figure 3.**
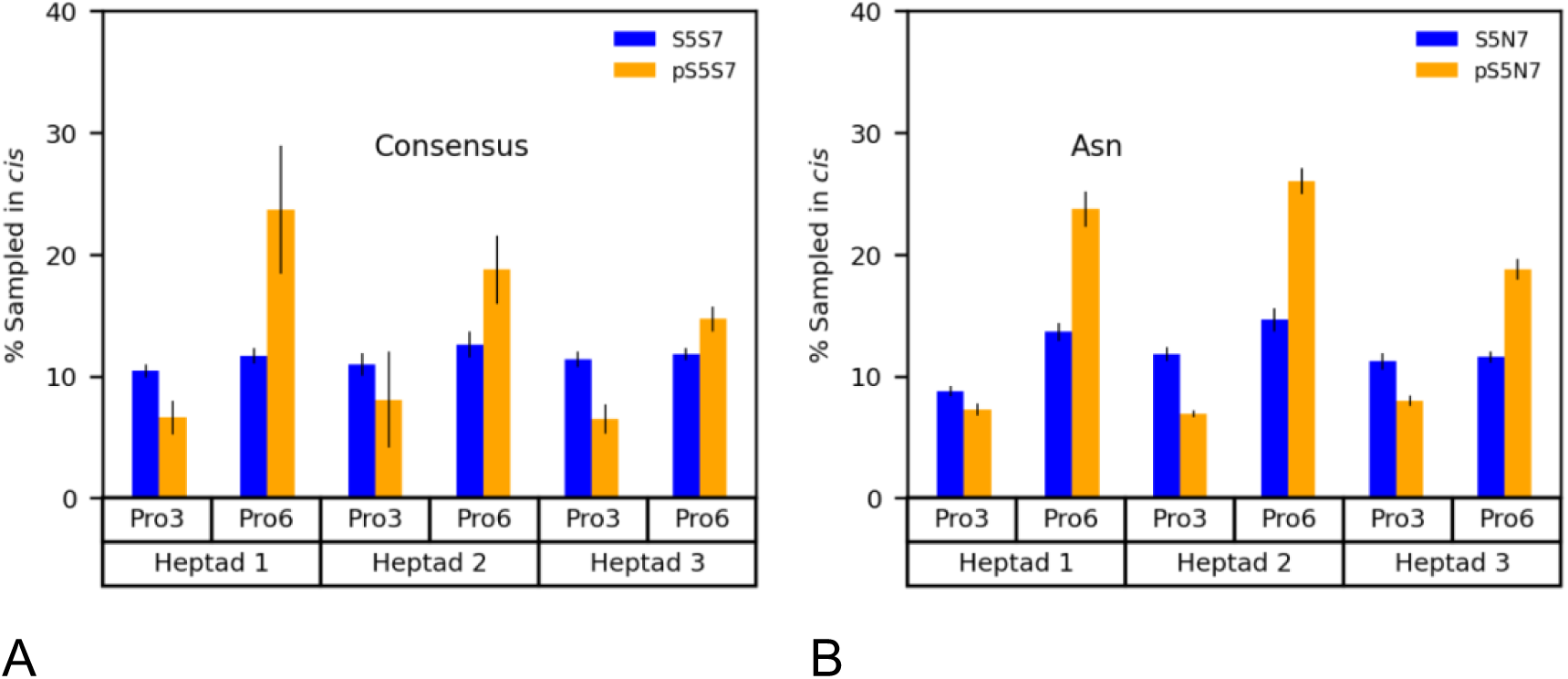
Percentage of simulations sampled in *cis* conformation at each Pro in the CTD. For pS5S7 and pS5N7 simulations, Ser5 in each heptad is phosphorylated (*A*) Consensus variant (*B*) Asn variant (Heptad 2 contains Asn at position 7)

**Figure 4.**
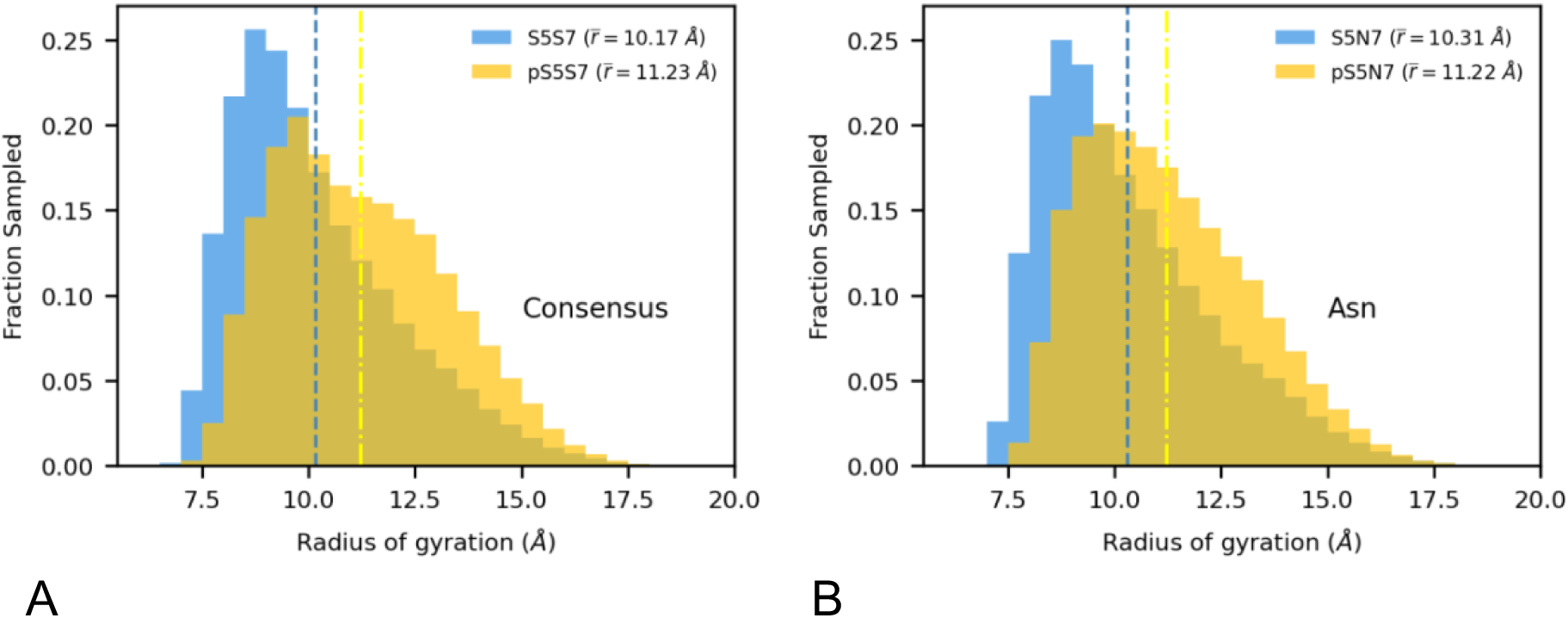
Histogram of the radius of gyration comparing unphosphorylated to phosphorylated ensembles, with blue, dashed line indicating average radius of gyration across unphosphorylated simulations and yellow, dash-dotted line indicating average radius of gyration across phosphorylated simulations for (*A*) consensus (*B*) Asn variant.

**Figure 5.**
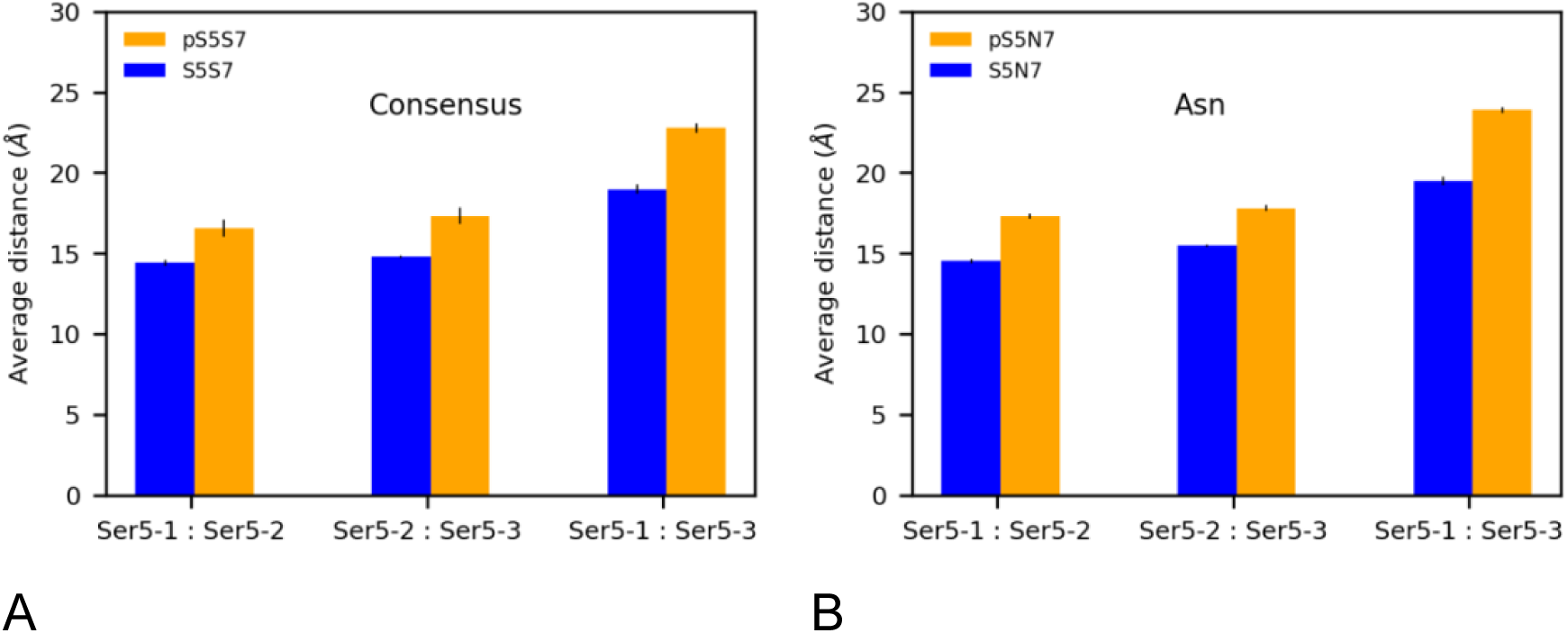
Distances between Ser5 side chain OG atom comparing the unphosphorylated to phosphorylated ensembles for (*A*) consensus (*B*) Asn variant. “-#” denotes the Heptad # in which the Ser5 residue analyzed belongs.

**Figure 6.**
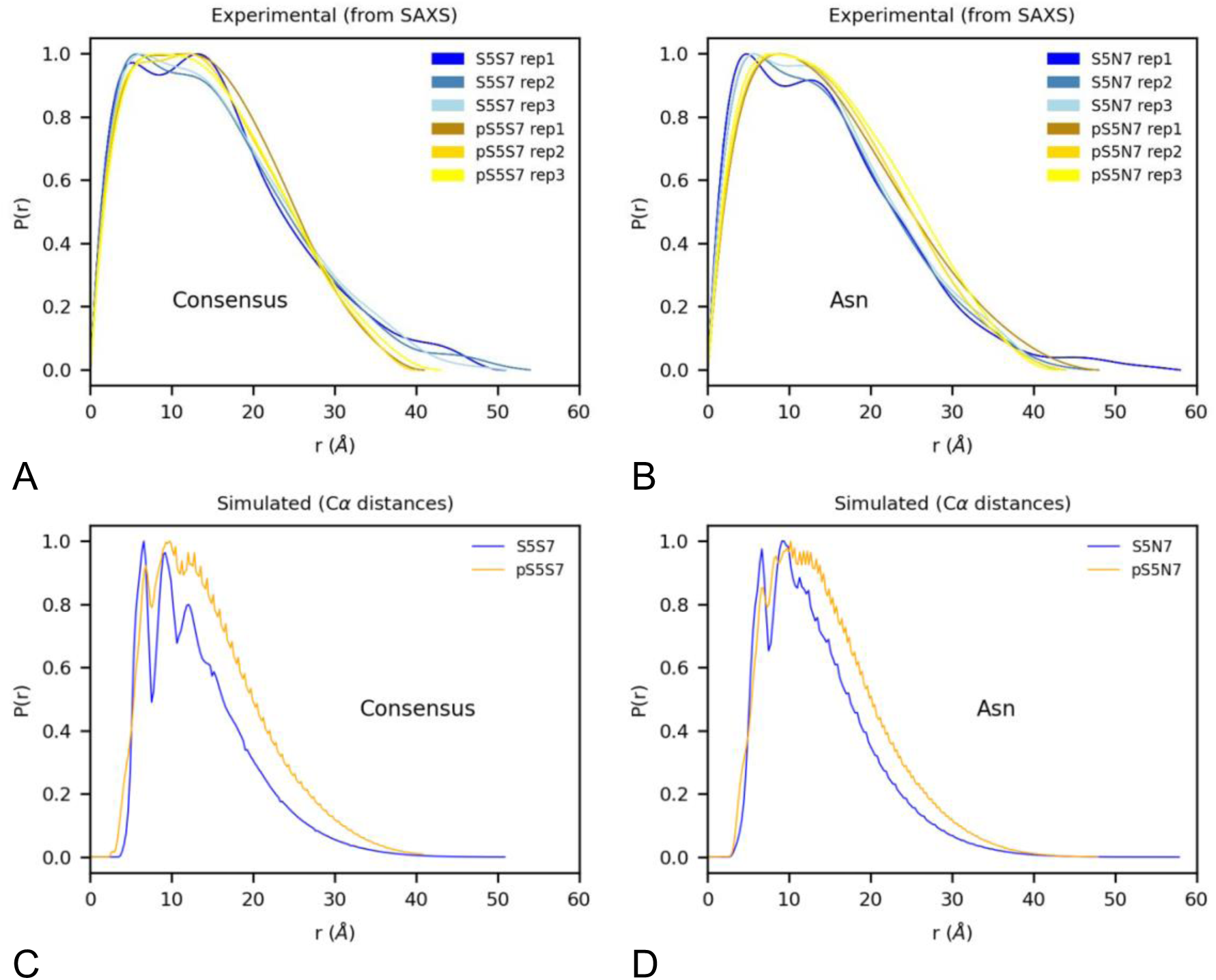
S**A**XS **data from experiments and comparison to simulated data. P(r) analysis from (A) experimental SAXS Consensus variant data (errorbars similar width to line plots), (B) experimental SAXS Asn variant data (errorbars similar width to line plots), see also Fig. S5), (C) P(r) Cα distances in simulated Consensus variant data, and (C) Cα distances in simulated Asn variant data. 3 replicates are displayed for each experimental construct in A and B. All P(r) data was normalized by dividing by the maximum value in the dataset.**

**Figure 7.**
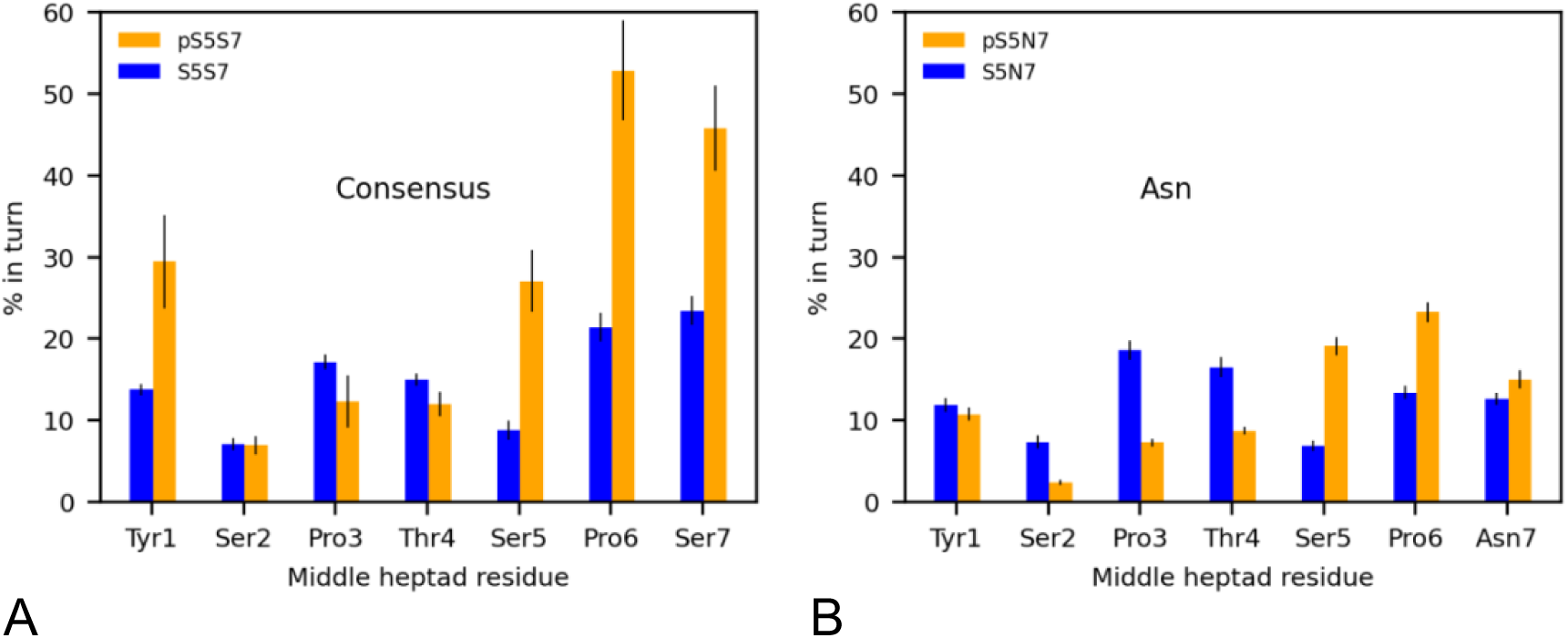
Percent turn character in CTD middle heptad residues across all unphosphorylated ensembles compared to phosphorylated ensembles for (*A*) consensus (*B*) Asn variant

**Figure 8.**
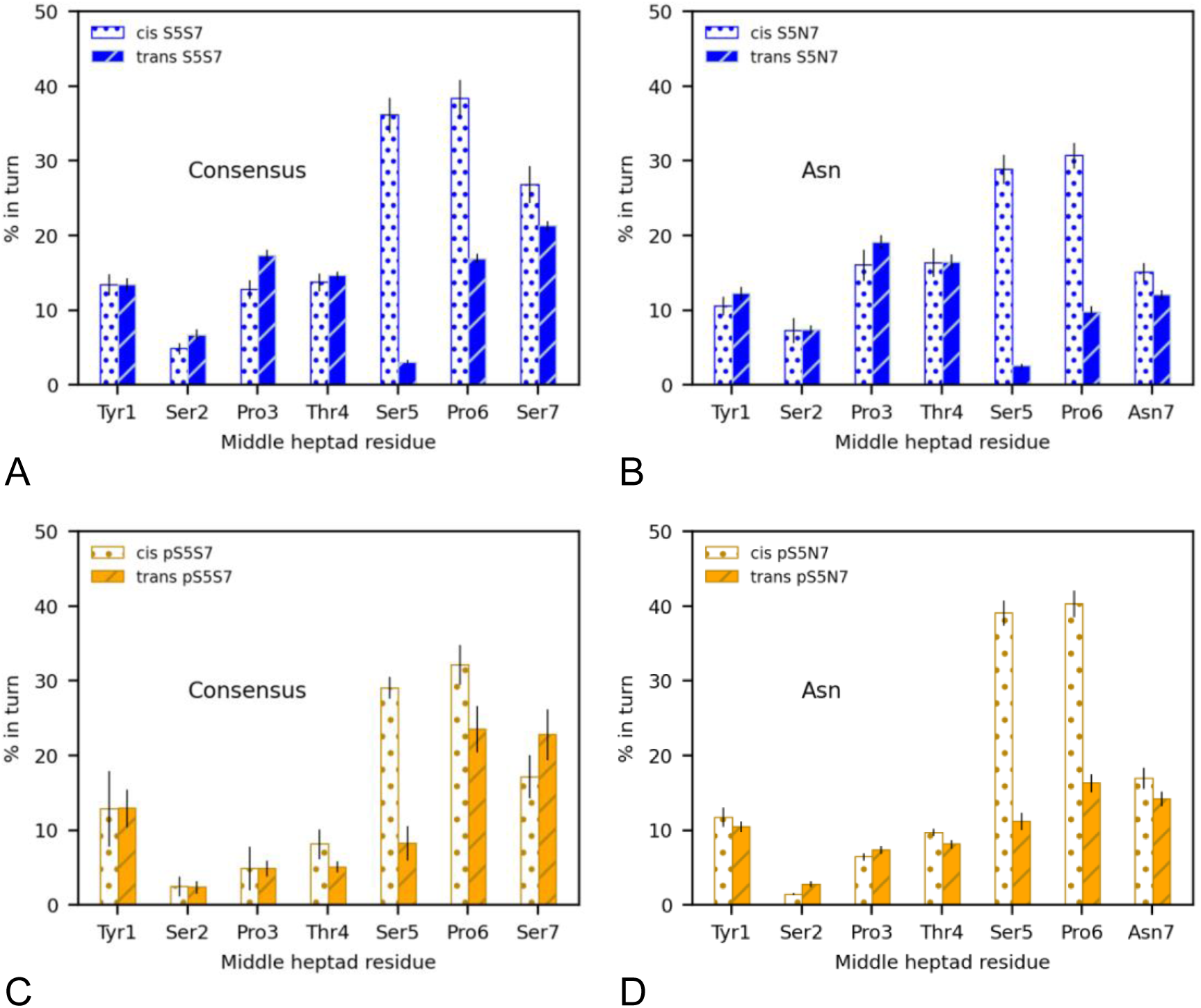
Percent turn character in CTD middle heptad residues with *cis*-Pro6 compared to *trans*-Pro6 (see also Fig. S6) in the (*A*) Unphosphorylated consensus, (*B*) Unphosphorylated Asn variant, (*C*) Phosphorylated consensus, (*D*) Phosphorylated Asn variant.

where *n* is the total number of simulations for that construct, and *w*_*i*_ is the weight for simulation *i*.

The weighted variance, *σ*^2^, is calculated by the following formula:

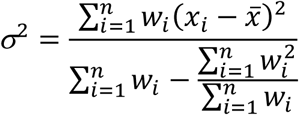

where *x*_*i*_ is the value of the mean for simulation *i*, and ^-^*x* is the average value of all the simulation means.

Then, the weighted standard error of the mean, *SE*, is calculated as follows:

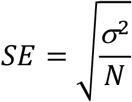

### Contact Map Analysis

To visualize how the global conformation of the CTD is altered by phosphorylation, multiple contact maps were created by utilizing the open-source Contact Map Explorer python package. An averaged contact map was created for each set of simulations, for every 100 frames of each simulation (every 10 ps), using the reweighted data. Each map consisted of binary 21 residue by 21 residue matrices, where 1 signifies a contact being present, and 0 does not. The cutoff distance for a contact was defined as 4.5 Å. After creating a list of all binary frames in a set of simulations, we applied the values from reweighting such that every binary frame was multiplied by its respective weight. The average of all reweighted frames was then taken for the final contact map. Residues that are adjacent to each other are not included in the final contact maps, as they are always going to be within 4.5 Å of one another.

The Scikit-learn *K*-means clustering algorithm was used on all simulation contact map data at once (taken every 100 frames, or 10 ps). To determine the optimal *k*-value, or the optimal number of clusters, the within-cluster sum of squares (WCSS) was plotted against *k*-values ranging from 1 to 30. Using this method, the ideal *k*-value should be located at the “elbow”, or the point at which the WCSS starts to decrease at a smaller rate, forming a bend in the plot. As this is a heuristic technique, and the elbow method only helps to approximate an appropriate k-value, we considered multiple *k*-values in the elbow region before selecting our final *k*-value of 11. With 11 clusters, we were able to visualize several clusters that were distinct from one another and showed differences in population when phosphorylated.

### Small-Angle X-Ray Scattering (SAXS)

Peptides S5S7, pS5S7, S5N7, pS5N7, with N-terminal acetylation and C-terminal amidation, were synthesized by GenScript. Peptides were resuspended in 10 mM sodium phosphate pH 7.5, 50 mM NaCl, 2 mM DTT, and purified by a Superdex S-100 Increase column (Cytiva). Concentrations were determined by the infrared spectroscopy-based Direct Detect Spectrometer (EMD Millipore). SAXS was subsequently performed in triplicate for each peptide at a concentration of 1 mg/mL, except for pS5N7, which was at a lower concentration due to its lower solubility.

SAXS experiments were performed on a Rigaku MM007 rotating anode X-ray source paired with the BioSAXS2000nano Kratky camera system. The setup includes OptiSAXS confocal max-flux optics, designed specifically for SAXS, and a highly sensitive HyPix-3000 Hybrid Photon Counting detector. The sample-to-detector distance was calibrated to 495.5 mm using silver behenate powder (The Gem Dugout, State College, PA). The momentum transfer scattering vector (*q*-space) range, defined as 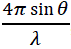, extended from qmin = 0.008 Å⁻¹ to qmax = 0.6 Å⁻¹ (where q = 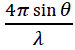 and 2θ is the scattering angle). The X-ray beam energy was set to 1.2 keV, with a Kratky block attenuation of 23% and a beam diameter of approximately 100 μm. For sample handling, protein samples were loaded into a quartz capillary flow cell using the Rigaku autosampler, with the sample stage maintained at room temperature. The entire X-ray flight path, including the beam stop, was held under vacuum conditions of < 1×10⁻³ torr to reduce air scattering. Automated data collection, including detailed cleaning cycles between samples, was managed by the Rigaku SAXSLAB software.

Data processing was completed using the Rigaku SAXSLAB software, with each dataset averaged over six ten-minute images across three replicates of both protein and buffer samples to confirm the absence of X-ray radiation damage. Consistency in SAXS data overlays verified no radiation decay or sample loss over the 60-minute collection period. Reference buffer subtraction was subsequently performed to isolate the protein’s raw SAXS scattering curve. The forward scattering *I*(0) and radius of gyration (*R*_g_) were calculated using the Guinier approximation, which assumes the intensity follows 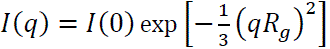 at very small angles (*q* < 1.3/*R*_g_). Data analysis included evaluating parameters such as radius of gyration, D_max_, Guinier fits, Kratky plots, and the pair distance distribution function with ATSAS software. The pairwise distance distribution function *P*(*r*) was generated using GNOM, following protocols for intrinsically disordered proteins (34, 35), where D_max_ was increased until *P*(*r*) smoothly approaches zero at larger *r*. For comparison across conditions and with simulations, *P*(*r*) curves were scaled so that the maximum value equals 1.

## Results & Discussion

### Proline isomerization state sampling using GaMD

GaMD simulations were used to sample the conformational ensemble of a three-heptad CTD peptide, both with and without phosphorylation at Ser5. The GaMD enhanced sampling method was required to overcome the free energy barrier between the *cis* and *trans* states of proline *ω* dihedral angle, allowing transitions between these conformations to happened frequently throughout each of the independent simulations (Fig. S1). Throughout these simulations, the CTD was highly disordered with little secondary structure (less than 3% helix and β-sheet for all residues). Conformations varied from compact to extended within a single 490-ns simulation (Fig. 2).

The reweighted ensemble generated from GaMD simulations was analyzed to determine how frequently the proline residues in the CTD peptides sample the *cis* conformation (Fig. 3). In the unphosphorylated ensembles, all proline residues adopted the *cis* conformation around 10-15% of the time, for both the consensus and Asn variant. We also see that phosphorylation at Ser5 increases the *cis* population at Pro6, especially for the Asn variant. Comparing these results to previous NMR experiments on a hyper-pSer5 11-heptad repeat segment of *Drosophila* CTD, there is agreement with the general trend that Ser5 phosphorylation increases the *cis*-Pro6 conformation, and more so for the Asn variant than the consensus peptide (4). Although the simulations show slightly higher *cis* population in the unphosphorylated ensemble (10%) than the NMR experiments (5%), this difference is on the order of the experimental error and could be due to a slight preference for the *cis* conformation by the force field used in our simulations, or the ability for the shorter CTD peptide (3 heptad repeats) to more easily adopt a *cis* conformation. Our observed increase in *cis* conformation for prolines in the *i*+1 position for the CTD contrasts with Ash1, which was not observed to have an overall change in *cis*-Pro occupancy with phosphorylation (22). Tau208-324 also still exhibited a dominant *trans* conformation at Pro232 with phosphorylation at Thr231(7). In the Ash1 sequence, looking specifically at phosSer followed by Pro-Asn (which is the sequence pattern we have in the Asn variant of the CTD), Martin *et al.* did see a small increase in the *cis* population on phosphorylation (increase from 7.2% to 11.6%) (22). Together, these results indicate that the specific sequence around the phosphorylation site is important and to generalize that phosphorylation increases *cis*-Pro in IDPs would be too broad. In fact, at Pro3 of the CTD heptads, we observed a decrease in the *cis* percentage upon Ser5 phosphorylation (Fig. 3), which we explain in a subsequent section.

### Ser5-phosphorylation leads to expansion of the peptide

Comparing the phosphorylated and unphosphorylated CTD peptide ensembles, we observe a slight increase in the average radius of gyration upon phosphorylation for both the consensus and Asn variant (Fig. 4). This effect is likely due to the charge repulsion between the negatively-charged phosphate groups, as the average distance between each of the Ser5 side chains increases significantly in the phosphorylated ensemble for both the consensus and Asn variant (Fig. 5). This effect has been observed before in both experiments and MD simulations (19), notably in a 2021 study by Jin and Grater, which included simulations of a longer CTD construct (20). The authors found that the radius of gyration was directly related to net charge, with neutral or negatively-charged IDPs expanding with phosphorylation, although this effect was more pronounced in simulations than in experimental data. The authors attributed this discrepancy to an inflation of electrostatic interactions in current protein molecular dynamics force fields.

We performed SAXS experiments (Fig. S2-S5, Table S1) and compared to the global conformation of the three-heptad repeat peptide in our simulations. SAXS data did not reveal a significant difference in the radius of gyration for the phosphorylated and unphosphorylated peptides (Fig. 6). However, the radii of gyration calculated from our simulations are in general agreement with those calculated from the SAXS data (Table 2), but the SAXS experiments are not able to detect the small change in expansion of the peptide seen computationally. The pair-wise distance distributions from simulation and SAXS experiments show a similar overall shape that changes little with phosphorylation (Fig. 6). Simulations show a slight increase in the population at the high end of the pair-wise distance distribution for the phosphorylated peptide, indicating fewer interactions between residues that are distal in sequence. However, this difference is not visible in the SAXS data pair-wise distribution plots. Previous studies have observed a difference in the hydration levels when an IDP is phosphorylated, which can mask changes in radius of gyration upon phosphorylation in the SAXS data (20). Given that the change we observe computationally is small, it is possible that this subtle difference would not be visible experimentally.

**Table 2.** Radius of gyration values calculated in Angstroms using Guinier and P(r) analysis on experimental SAXS data and radius of gyration histograms from simulated data.

| Construct | Guinier | SAXS $P(r)$ | Simulated |
| --- | --- | --- | --- |
| S5S7 | $11.64 \pm 0.08$ | $12.83 \pm 0.48$ | $10.17 \pm 1.94$ |
| pS5S7 | $10.86 \pm 0.07$ | $11.85 \pm 0.61$ | $11.23 \pm 1.99$ |
| S5N7 | $11.25 \pm 0.08$ | $12.13 \pm 0.51$ | $10.31 \pm 2.00$ |
| pS5N7 | $11.27 \pm 0.15$ | $12.46 \pm 0.25$ | $11.22 \pm 1.94$ |

### Ser5 phosphorylation promotes local turn conformations

The *cis* conformation of the proline *ω* dihedral angle is associated with a kink in the protein backbone, which is associated with turn and bend secondary structures (10, 13). Our simulations therefore show an increase in turn conformations around Pro6 with Ser5 phosphorylation from 21% to 53% for the consensus peptide and from 13% to 23% for the Asn variant (Fig. 7), focusing on the conformation of the middle heptad to minimize any effects of being near the chain terminus. For both sequences, we also observe a drop in the turn conformation around Pro3 when Ser5 is phosphorylated, consistent with the decrease in *cis* conformations at Pro3. Fig. 8 demonstrates that there is a clear correlation between the conformation of the Pro6 *ω* dihedral angle and the turn conformation of the peptide backbone. Interestingly, the increase in turn conformations on phosphorylation of the Asn variant cannot be accounted for purely through the increase in *cis* conformations, as even the amount of turn among the *trans* conformations increase after phosphorylation (Fig. S6). Additionally, even among the structures with Pro6 in a turn conformation, the percentage of time in *cis* increases with phosphorylation (Fig. S6). This emphasizes that both turn conformations and *cis* conformations, which are characterized by an overall change in direction of the peptide backbone, are stabilized by phosphorylation at Ser5.

### Increased proline *cis* conformation with Ser5 phosphorylation counteracts expansion due to negative charge

The repulsion between the negatively-charged Ser5 phosphate groups and the increase in *cis*-Pro6 and turn conformations have opposite effects on the global properties of the peptide ensembles. While the electrostatic repulsion leads to expansion, this is moderated by the compacting effect of the increased *cis* and turn conformations for Pro6, especially for the Asn variant. This can be seen by comparing the radius of gyration for the subset of the ensemble with Pro6 in turn to Pro6 not in turn (Fig. 9 *A*-*D*). When Pro6 is not in turn, we see that the phosphorylated ensemble samples many more of the expanded conformations with a radius of gyration greater than 13 Å, especially for the consensus variant. The structures with Pro6 in a turn conformation have a smaller radius of gyration, making the expansion from phosphorylation less pronounced. Although there is an increase in the turn conformation at Pro6 with phosphorylation for the Asn variant, there is a decrease at Pro3, and across the entire peptide, phosphorylation does not affect the total amount of turn structure present (Fig. S7 *A*). However, there is an increase in the average number of prolines in *cis* with phosphorylation (Fig. 3), and there is a clear correlation between the radius of gyration and the total number of the prolines in the sequence that are in cis for both the consensus and Asn variant CTD peptides (Fig. 9 *E*-*F*).

**Figure 9.**
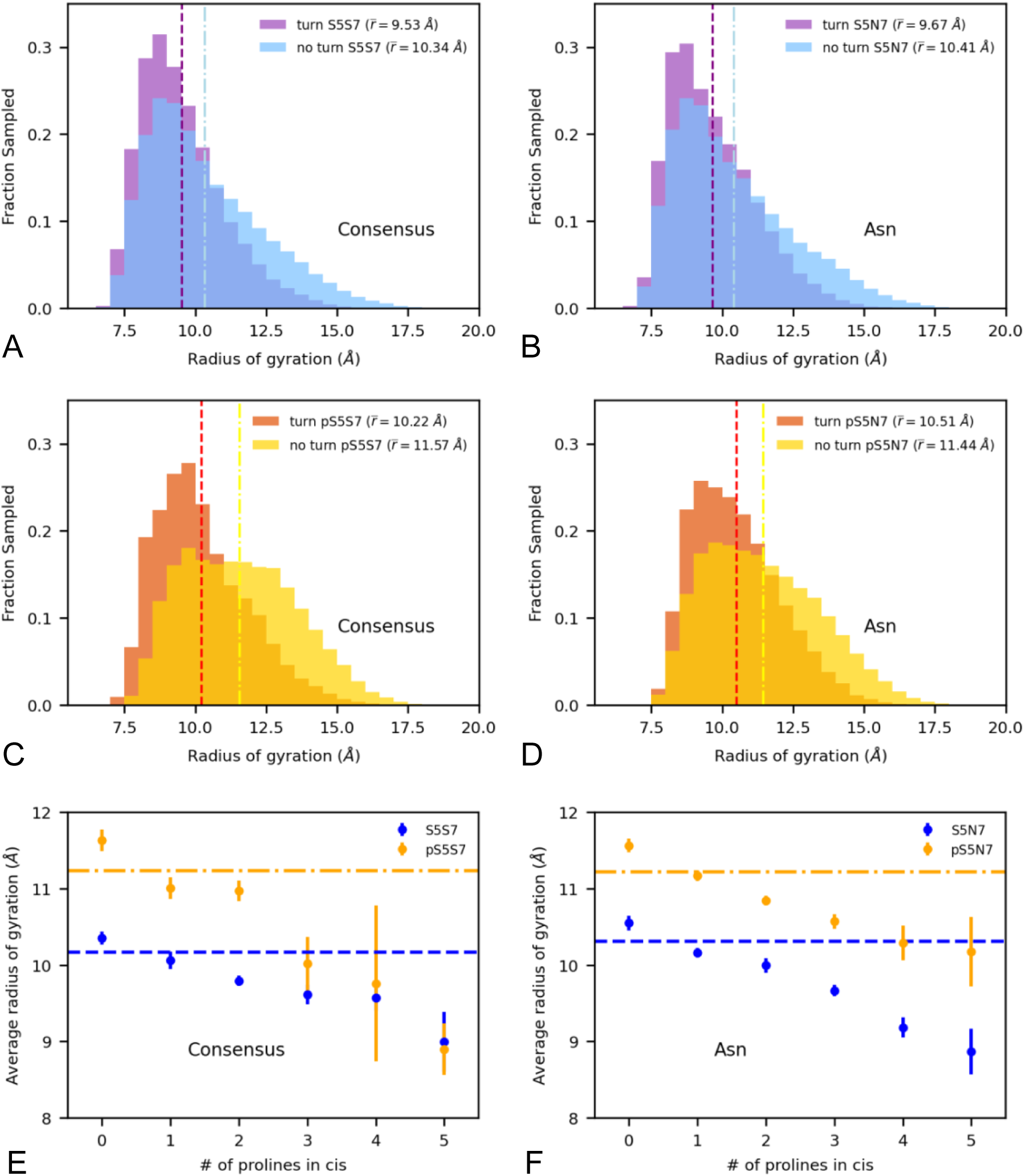
Effect of proline conformation on radius of gyration. Histogram of the radius of gyration comparing when Pro6 is in turn or not in turn for (*A*-*B*) consensus (*C*-*D*) Asn variant. Dashed lines indicate average radius of gyration across all frames where Pro6 is in turn, and dash-dotted lines indicate average radius of gyration across all frames where Pro 6 is not in turn. Scatter plot of average radius of gyration vs number of proline residues in the *cis* conformation for (*E*) consensus and (*F*) Asn variant. Blue, dashed lines indicate average radius of gyration across unphosphorylated simulations, and orange, dash-dotted lines indicate average radius of gyration across phosphorylated simulations, regardless of how many proline residues are in the *cis* conformation.

This result, that the cis proline conformations are associated with a more compact ensemble, has been observed previously in MD simulations of proline-rich IDPs (16, 17). These cis conformations are increased with phosphorylation at the Pro6 position in each heptad, multiplying the compacting effect by the number of heptads. We also see that phosphorylation has an expanding effect for a fixed number of prolines in *cis*, indicating that the expanding and compacting effects are independent and competing in the phosphorylated ensemble (Fig. 9 *E*-*F*).

Previous SAXS experiments on the hyper-pSer5 11-heptad CTD showed a minimal effect of phosphorylation on radius of gyration and pair-wise distance distributions (4). This may be attributed to the competing effect of expansion from electrostatic repulsion and compaction from increased *cis*-Pro6 conformations. Previous MD simulations of this 82-residue CTD construct showed a larger change in radius of gyration of 14 ± 6 Å, which the authors attributed to an inflation of electrostatic interactions by common MD force fields used with IDPs (20). We propose that at least part of this discrepancy may be because the simulations performed previously with standard MD were not able to sample *cis* conformational states of the proline residues in the protein. Our simulations on a shorter CTD construct show that if only *trans* conformations of proline were included the peptide would show greater expansion with phosphorylation, as can be seen by looking at Fig. 9 *E*-*F* for zero prolines in *cis*. This effect would be larger with a longer 11-heptad peptide which has more proline residues that can sample the *cis* conformation.

In order to gain more insight into the global conformational effect of phosphorylation, we constructed intramolecular contact maps that display the frequency of interactions between CTD residues (Fig. 8H). When comparing the contact maps for the phosphorylated and unphosphorylated peptide, we can see that certain local contacts (near the diagonal) increase with Ser5 phosphorylation, while others decrease (Fig. 10). Long-range interactions between residues distant in sequence generally decrease with Ser5 phosphorylation. The exception is the contacts between Ser5 residues on different heptads that increase with phosphorylation at these residues in the consensus peptide. However, despite an increase in those specific serine residues, the average distance between the serine side chains still increases with phosphorylation for the consensus peptide (Fig. 5).

**Figure 10.**
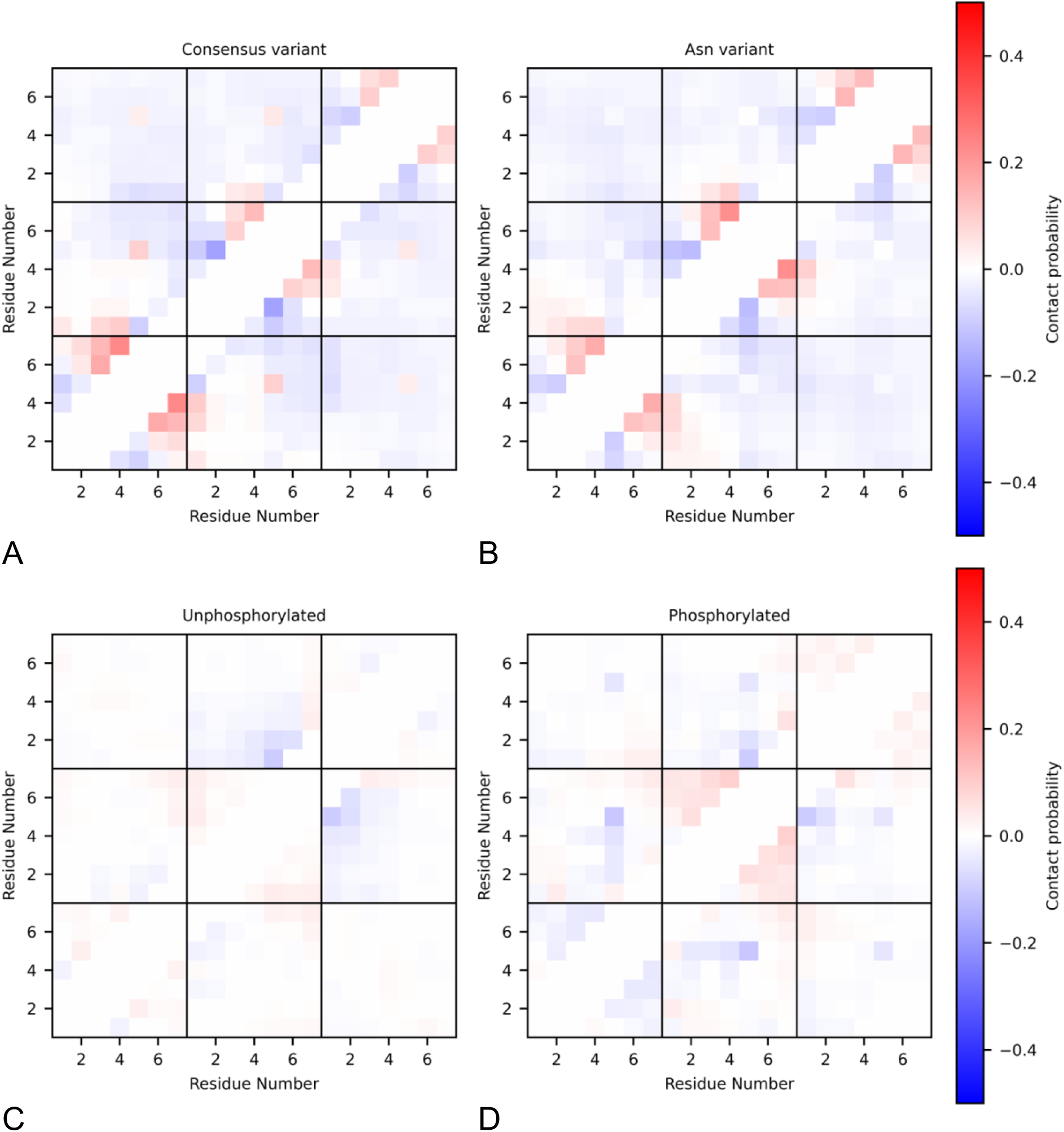
Contact difference maps of the probability of residues in contact subtracted for the following ensembles (see also Fig. S8): (*A*) Consensus, phosphorylated minus unphosphorylated ensembles (*B*) Asn variant, phosphorylated minus unphosphorylated ensembles (*C*) Unphosphorylated, Asn minus consensus variant (*D*) Phosphorylated, Asn minus consensus variant

### Phosphorylation shifts the population of local structural features

To gain insight into the particular conformations sampled in the CTD peptide ensembles and the differences in the local contacts, we performed k-means clustering on the combined contact map data from all of our simulations combined. We used the within-cluster sum of squares (WCSS) to measure the deviation of individual conformations from their cluster mean and determine the appropriate number of clusters. We plotted this value against the number of clusters and identified an “elbow” in the plot at six clusters and at eleven clusters, indicating that the improvement levels off after that these *k*-values (Fig. S9). We chose to use eleven clusters because this number included several clusters that showed differences in population when phosphorylated. The contact maps associated with each of the eleven clusters are provided in Fig. S9 and additional data on each cluster is in SI Table O.

Here we focus on six of the clusters that provide insight into the role of phosphorylation and heptad sequence on local conformation (Fig. 11). Cluster 5 has the highest cluster population, representing structures where no contacts occurred. As expected, it is the cluster with the largest radius of gyration (12.2 Å), and it has a slightly higher occupancy in the phosphorylated ensembles due to the repulsion between the negative phosphate groups, consistent with our radius of gyration histograms (Fig. 4). Repulsion between phosphorylated serine residues also explains the decrease in occupancy of Cluster 3 in the phosphorylated ensembles. Cluster 3 is the most compact cluster of those with significant occupancy in the reweighted ensemble, with a radius of gyration of 8.9 Å. It is characterized by a turn in the middle heptad and interaction between the Tyr1 residues in the outer heptads (Fig. 11). The interaction between the tyrosine residues is characterized by a hydrogen bond in 6% of the structures in Cluster 3.

**Figure 11.**
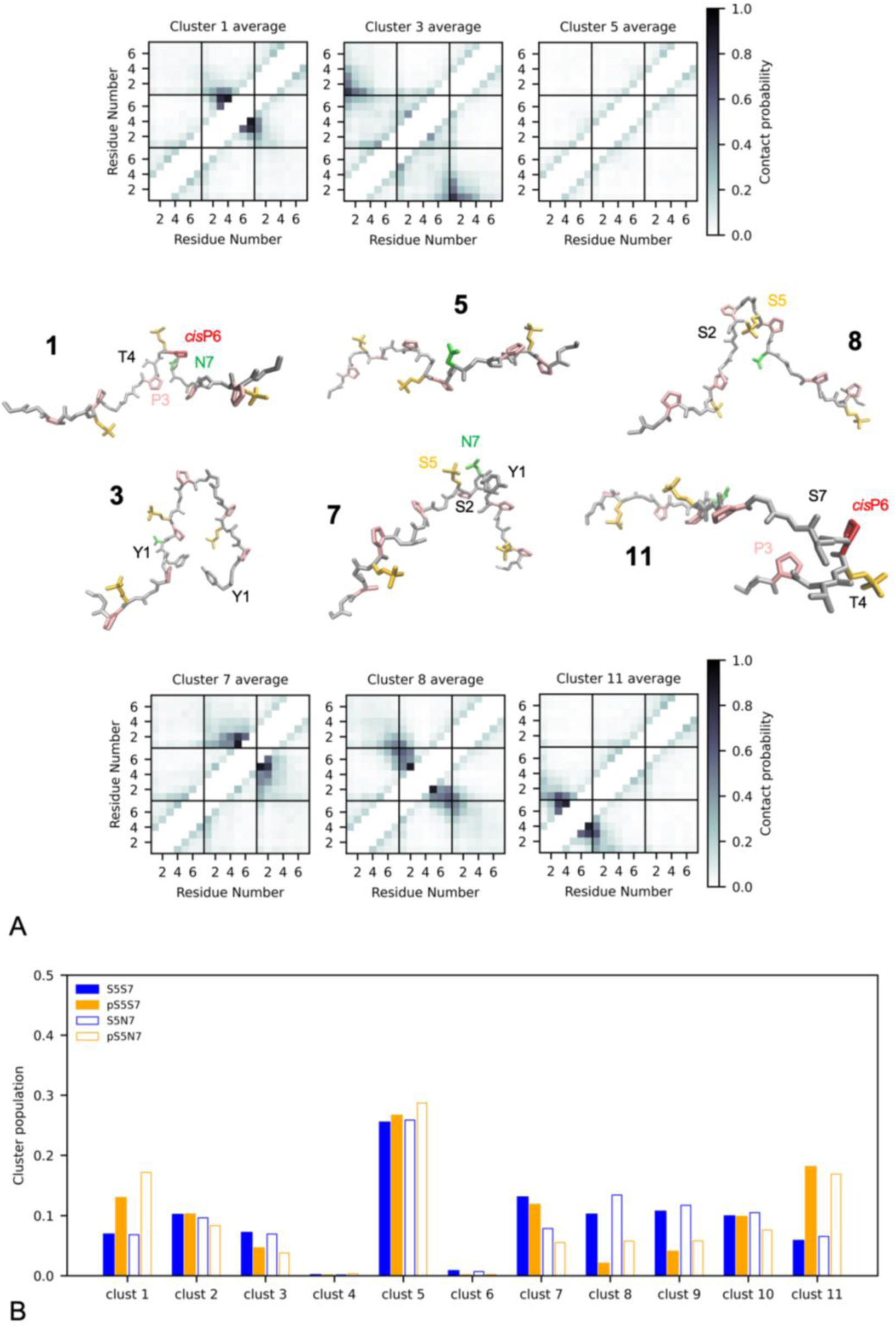
Clustering analysis (*A*) Cluster contact maps with centroid structures (*B*) Population comparison of each cluster for the different peptides (see also Fig. S9).

Clusters 8 and 9 are notable because they represent structures that are destabilized by phosphorylation. Cluster 8 is characterized by a contact between Ser2 and Ser5 in the middle heptad (Fig. 11), with a hydrogen bond forming between the two serines in 21% of the structures in that cluster. This also corresponds to a turn conformation at Pro3 between the two serine residues and is associated with a higher chance of Pro3 being in a *cis* conformation (45% of the time in turn and 15% of the time in *cis*). This turn conformation in the middle heptad also means that Cluster 8 is one of the more compact clusters with a radius of gyration of 9.0 Å, and tends to keep the Ser5 side chains closer together. Cluster 9 represents the same interaction between Ser2 and Ser5 as Cluster 8, but in the third heptad rather than the middle heptad (Fig. S9).

Phosphorylation likely disrupts the interaction between Ser2 and Ser5 as the large negatively-charged phosphate group takes up more space, making it geometrically harder to form this interaction with the bulky side chain. Phosphorylated Ser5 residues in Cluster 8 are less solvent exposed than on average in the phosphorylated ensembles (157 Å^2^ solvent exposed surface area compared to an average value of 170 Å^2^). Repulsion between phosSer5 residues may also contribute to the decrease in this conformation for the phosphorylated peptides, since this more compact structure brings the Ser5 residues closer together than average. The decrease in this conformation also explains why we see a decrease in both turn and *cis* conformation at Pro3 with phosphorylation.

Cluster 7 also represents an interaction with Ser5 of the middle heptad that is destabilized by phosphorylation. In this case, Ser5 interacts with residue 7, which is serine in the consensus sequence and asparagine in the Asn variant (Fig. 11). Ser5 is also in contact with Tyr1 of the following heptad. Interestingly, this conformation is associated with a turn conformation at Pro6 (48%), but not with a *cis* conformation of the Pro6 peptide bond (12%). In contrast, Cluster 1 is highly associated with a turn and *cis* conformation at Pro6 of the middle heptad (49% in turn and 53%). The *cis* conformation of the Pro6 peptide bond tends to point the Ser5 and Ser/Asn7 side chains away from each other, alleviating steric clashes (Fig. 12). This explains why Cluster 7 is less populated in the phosphorylated ensembles while Cluster 1 is more populated. This effect is also more pronounced for the Asn variant, which populates Cluster 7 less than the consensus and Cluster 1 more than the consensus, due to the bulkier branched asparagine side chain. Cluster 7 is the conformation where Ser5 is the least solvent exposed (54 Å^2^ for Ser5, 141 Å^2^ for phosSer5), while Cluster 1 has the most solvent exposed Ser5 residue (98 Å^2^ for Ser5, 183 Å^2^ for phosSer5).

**Figure 12.**
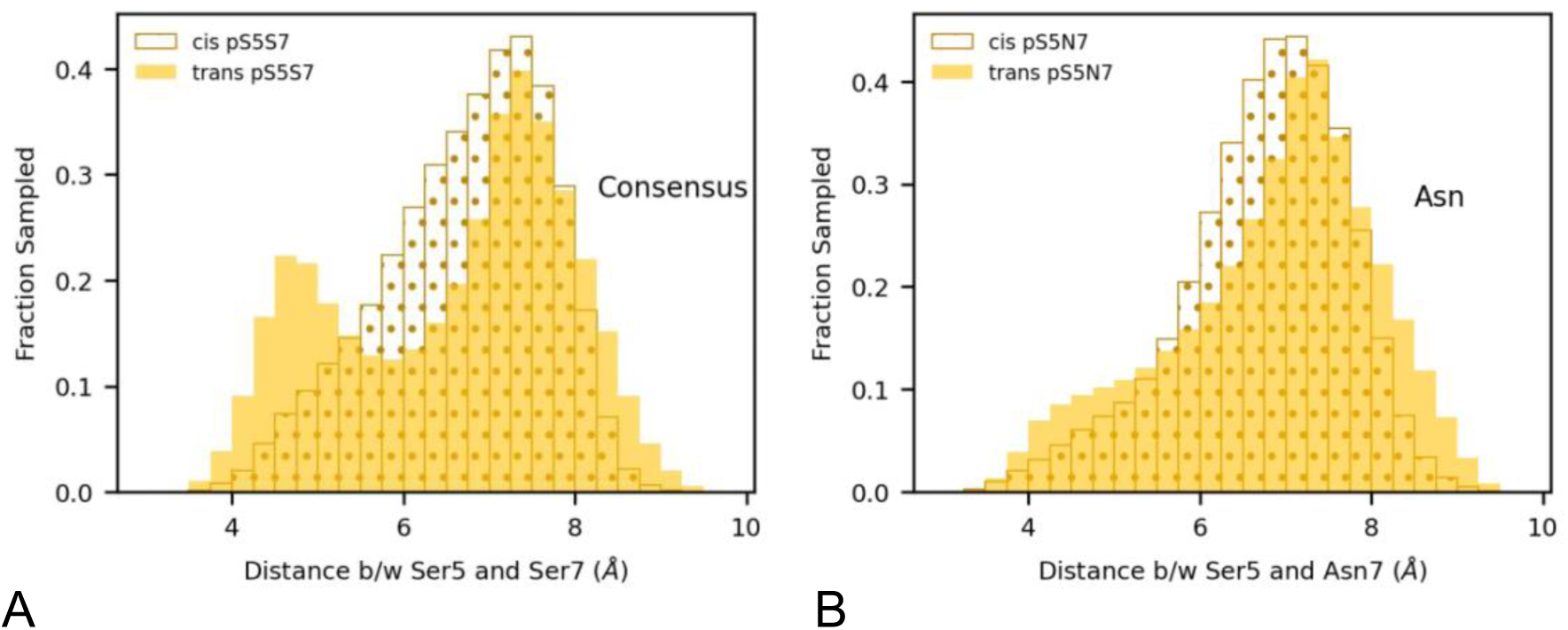
Histograms of the distances between the beta carbons for phosSer5 and Ser/Asn7 of the middle heptad for the (A) consensus peptide and (*B*) Asn variant comparing conformations when Pro6 of the middle heptad is in *cis* or *trans*.

Cluster 1 also tends to involve an interaction between residue 7 and Thr4 (in 15% of the cluster, this interaction is stabilized by a hydrogen bond), which could stabilize the *cis*-Pro6 conformation, especially in the Asn variant. Cluster 11 represents a similar *cis*-Pro6 and turn conformation to Cluster 1, but in the first heptad rather than the middle heptad. In this case, the population of Cluster 11 is higher in the phosphorylated ensembles but shows no difference between the consensus and Asn variant since residue 7 of the first heptad is serine in both constructs.

Overall, phosphorylation at Ser5 has two major effects on the local CTD conformation. First, it repels the other phosSer5 residues, favoring more expanded conformations over more compact conformations. Second, it interacts more specifically with the residue in position 7 of the heptad, favoring conformations where the large phosphate group can point away from the residue 7 side chain. Fig. 12 *A* shows how the dihedral angle of the Pro6 peptide bond affects how close the phosSer5 side chain is to the residue 7 side chain by plotting the distance between the β-carbon atoms. When the consensus peptide Pro6 is in *trans*, we observe two populations, one with the phosSer5 and Ser7 side chains pointed in the same direction as in Cluster 7 (β-carbon distances below 6 Å) and one with the side chains pointed away from each other (β-carbon distances above 6 Å). When it is in *cis*, we see that the side chains of phosSer5 and Ser7 are more likely to point away from each other (77% above 6 Å, versus 67% when in *trans*). In the Asn variant, we see a subtler effect of the *cis* conformation of Pro6, resulting in the phosSer5 and Asn7 side chains pointing away (80% above 6 Å, versus 77% when in *trans*). For the Asn variant, we also see that even in the *trans* conformation, the side chains are more likely to point away from each other than for the consensus peptide as a result of the larger asparagine side chain. Because the peptide is more flexible with Pro6 in the *trans* conformation than in the *cis* conformation, there is no change in the average distance between the β-carbon atoms, but the histogram reveals that the structures with the shortest distances and largest steric interaction between the side chains are reduced in the *cis* conformation.

To see if the steric repulsion between the side chains is related to solvent exposure, we also compared the solvent-accessible surface area (SASA) for phosSer5 when Pro6 is in *cis* or *trans*. As with the β-carbon atom distance, we see more occupancy of low SASA values when Pro6 is in *trans*. However, when Pro6 is in *cis*, the phosSer5 side chain is held in a more solvent-exposed conformation away from the residue 7 side chain, so the SASA values are shifted higher (Fig. S10). When Ser5 is phosphorylated, it becomes more favorable that the phosphate group be solvent exposed, which drives the shift away from Cluster 7 and toward Cluster 1, where Pro6 is in *cis*. This holds the Ser5 side chain away from the residue 7 side chain, particularly for the Asn variant. This tendency for phosphorylation to favor solvent-exposed conformations is similar to observations from simulations of Sic1 and Ash1, which found that phosphorylation leads to more hydrogen bonds with water and more ordered water molecules around the phosphorylation sites (23).

## Conclusions

By using GaMD simulations to observe a greater sampling of *cis*-Pro isomerization states, we can identify characteristics inherent to these states that cannot be observed through experiment, or through traditional MD simulations. In our simulations, we see that phosphorylation of Ser5 promotes *cis*-Pro conformations, especially for the Asn variant, as is consistent with previous NMR results (4). We also see that there is little change in the overall radius of gyration with phosphorylation, which is consistent with SAXS data. This agreement with experimental results gives us confidence that we can use the atomic resolution structures from our simulations to learn about local structural changes in the CTD peptide ensemble that result from phosphorylation. We find that Ser5 phosphorylation has competing effects on the CTD structure, promoting extended conformations that increase the average distance between Ser5 residues and increasing compact conformations associated with *cis*-Pro6. While the extended conformations are a result of the repulsive negative charge of the phosphate group, the increase in *cis*-Pro6 conformations seems to be a result of the large, hydrophilic side chain of the phosphorylated serine, which is more stable in solvent-exposed conformations. While there is not a large change in the overall radius of the CTD when phosphorylated, there are changes in local structure and shifts in which conformations are sampled frequently. These structural features within the disordered CTD ensemble may be important for transcriptional regulation. Our results also highlight how sequence-specific the conformational effects of phosphorylation are, raising the possibility that variations in the heptad sequence may play an important functional role.

## Data Availability

Data that support the findings of this study are available from the corresponding author upon reasonable request.

## Author Contributions

K.A.B and S.A.S. designed the research and lead the project. W.B. and R.B.E. ran the molecular dynamics simulations. M.R.C., W.P.B., K.A.S., O.C.O., R.B.E., and K.A.B. performed computational analysis. W.C. and S.M.D. performed the experiments. W.C. analyzed the experimental data. M.R.C., K.A.S., and W.P.B. created the figures. K.A.B. and M.R.C. wrote the manuscript. M.R.C., W.C, W.P.B., K.A.S., R.B.E., O.C.O., S.A.S., and K.A.B. edited the manuscript.

## Supplemental Information

Document S1. Supplemental analysis, Figures S1-S10, and Table S1

## Acknowledgements

Thank you to Michael Donnelly for computational support. K.A.B. thanks the MERCURY Consortium for mentoring support. The co-authors acknowledge the X-Ray Crystallography and Scattering Facility, Huck Institutes of the Life Sciences, The Pennsylvania State University, University Park, PA (RRID:SCR_024464) for use of the biomolecular X-ray scattering equipment and Dr. Neela Yennawar for collecting the data shown in Fig. 6.

This work was supported by National Science Foundation grant MCB-1932730 and MCB-2428716 to S.A.S., and MCB-2324974 to K.A.B. Support was provided by the Skidmore College Faculty Student Summer Research Program to K.A.S. and O.C.O. W.C. acknowledges support from the Eberly Postdoctoral Research Fellowship.

