## Supplemental Material for "RNA polymerase II CTD Ser5 phosphorylation induces competing effects of expansion and compaction"

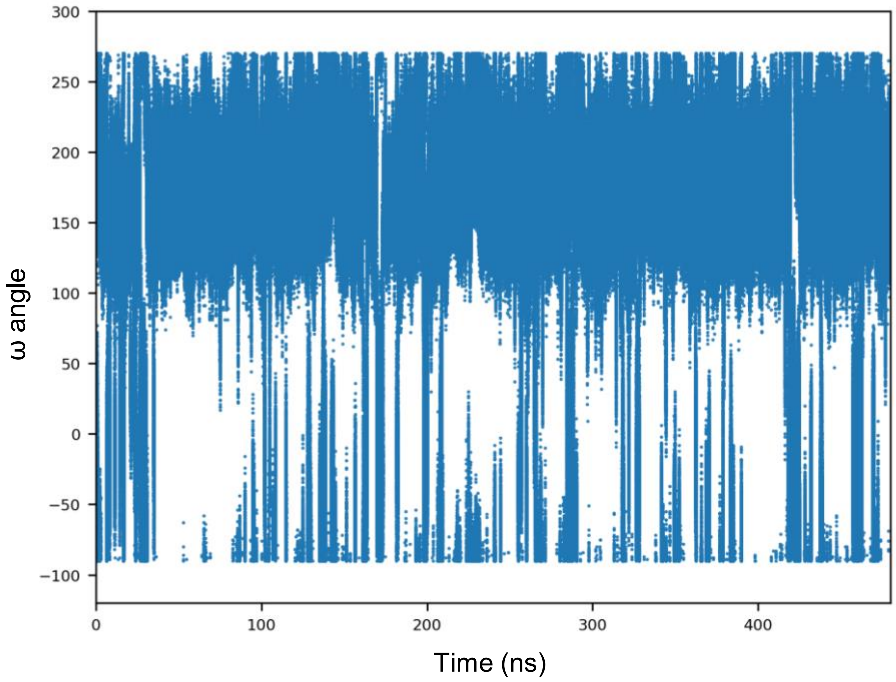


**Figure S1. Pro6 ω dihedral angle over the course of a single simulation.** The average isomerization frequency is ~10.7 flips/ns.

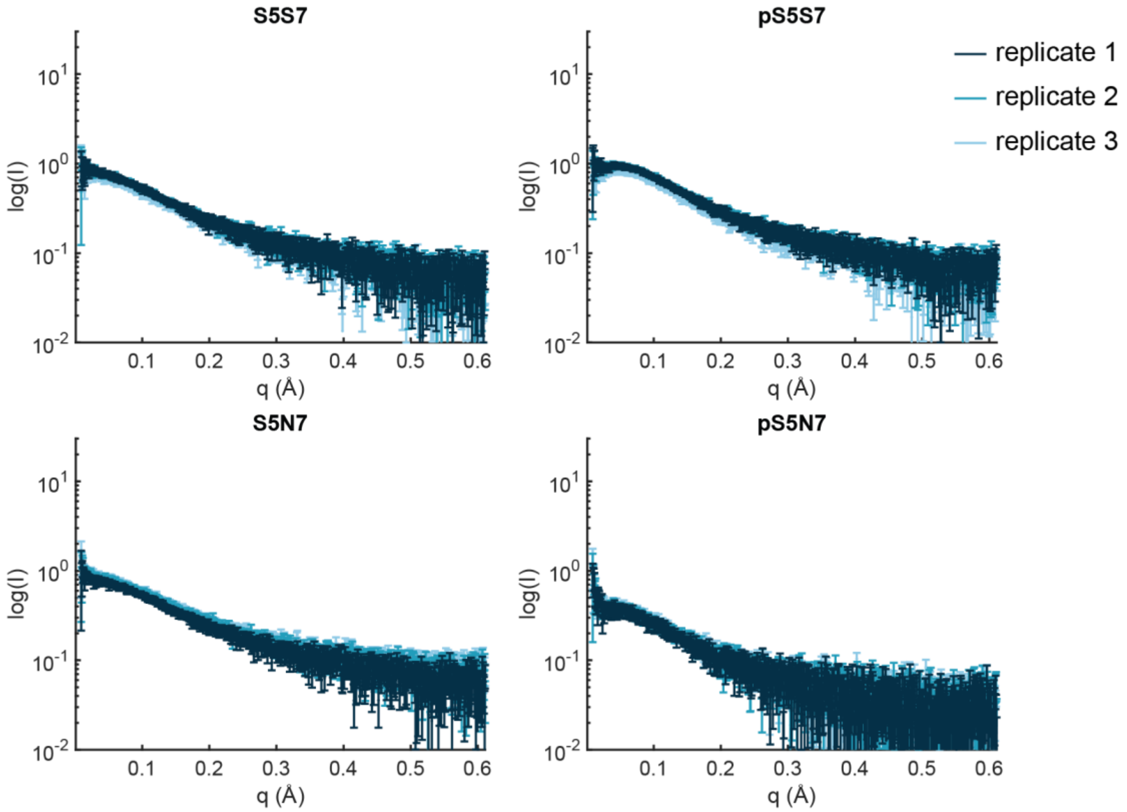


**Figure S2. Small-angle x-ray scattering (SAXS) data for CTD peptides.** Three technical replicates were performed for each peptide in 10 mM sodium phosphate pH 7.5, 50 mM NaCl, and 2 mM DTT at a concentration of 1 mg/mL, except for pS5N7 due to solubility issues. Scattering intensity at q = 0 suggests that the concentration of pS5N7 was about half of those of other three peptides.

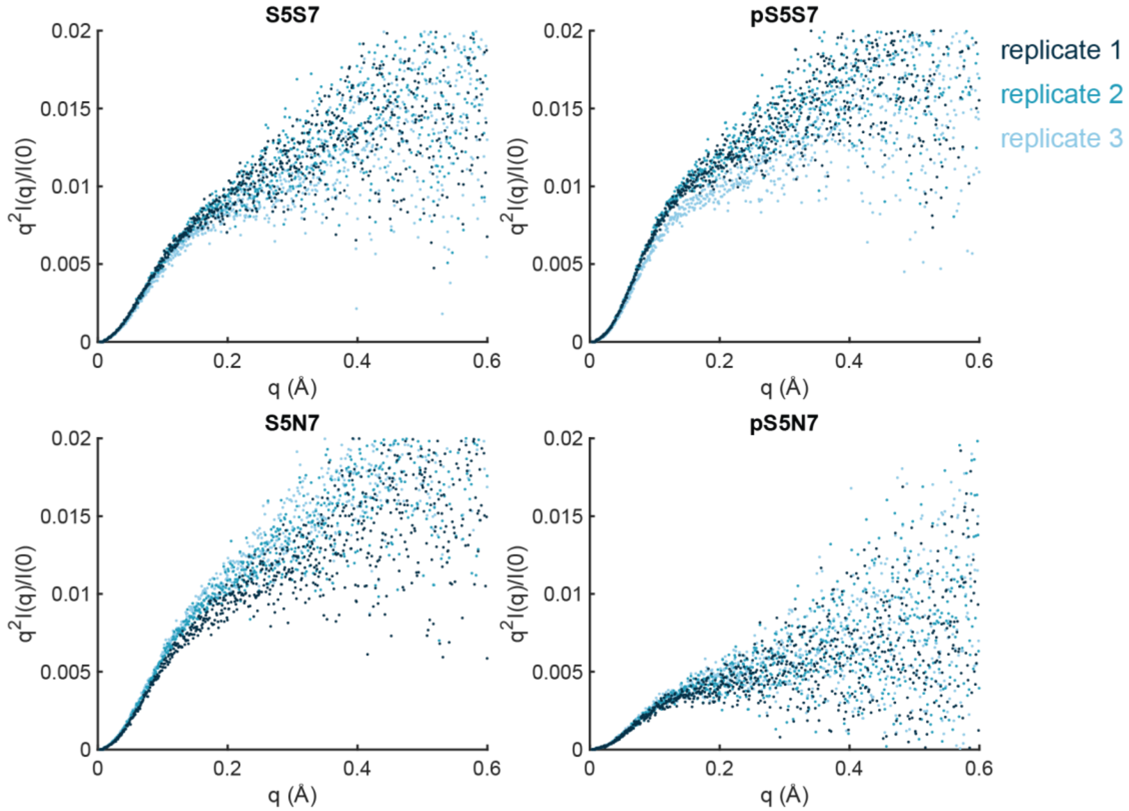


**Figure S3. SAXS Kratky plots for CTD peptides.** All peptides display a plateau for intrinsically higher q values, reflecting the extended, flexible nature of the polypeptide chain.

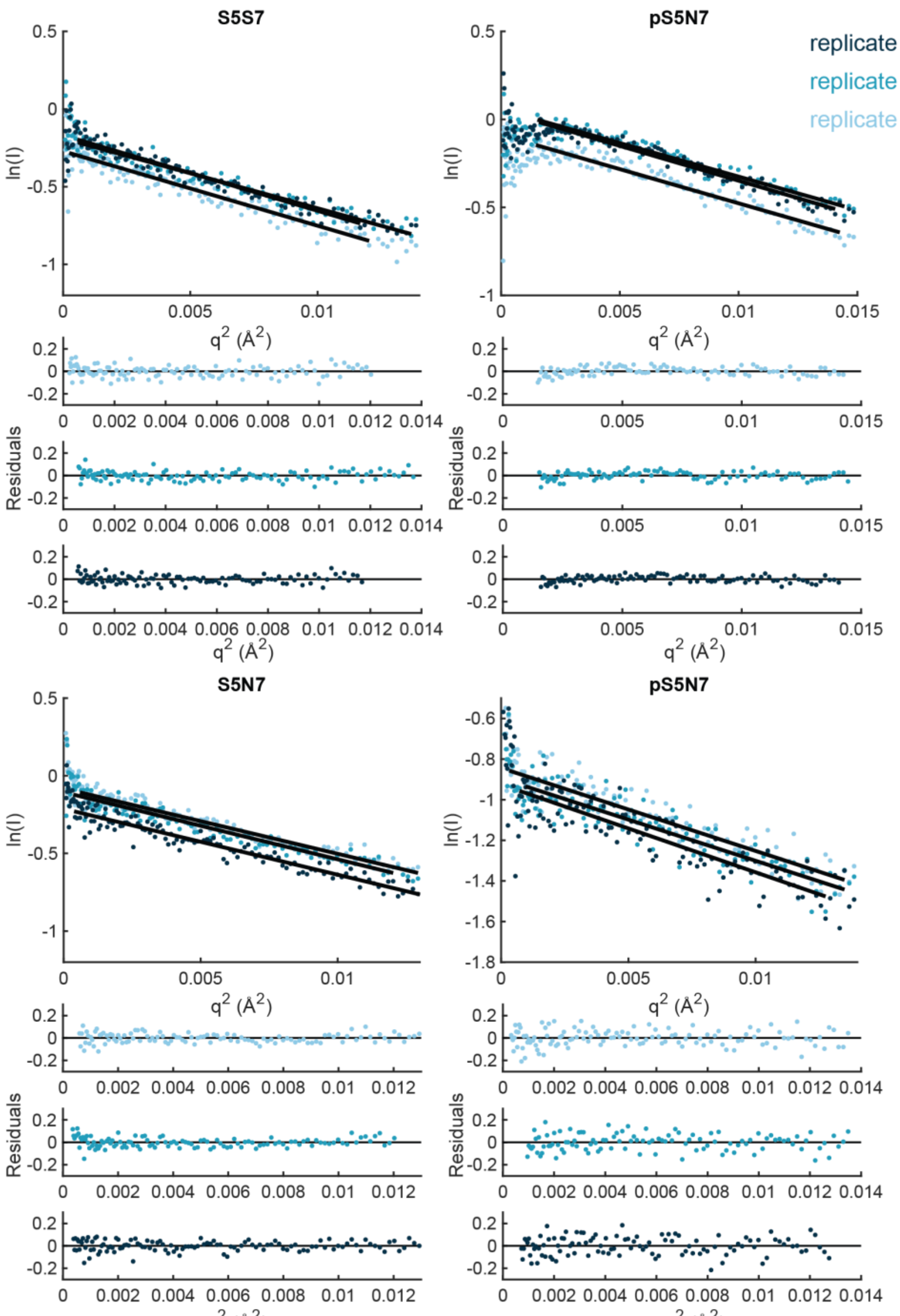


**Figure S4. SAXS Guinier analysis for CTD peptides.** Guineir fitting and residuals for CTD peptides. The resulting radii of gyration (Å) from fitting are: 11.85 ± 0.17, 11.05 ± 0.12, 12.02 ± 0.14 for S5S7, 10.88 ± 0.14, 10.75 ± 0.13, 10.95 ± 0.11 for pS5S7, 11.24 ± 0.14, 11.24 ± 0.15, 11.27 ± 0.14 for S5N7, and 11.18 ± 0.23, 11.16 ± 0.24, 11.48 ± 0.29 for pS5N7.

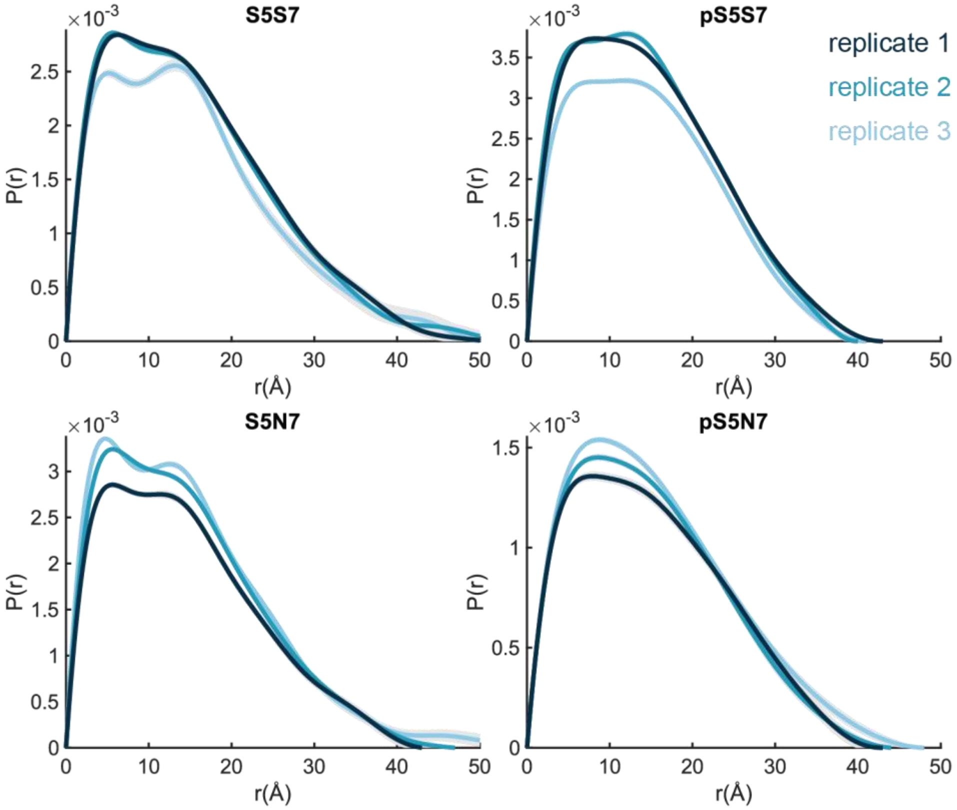


**Figure S5. SAXS pair-wise distance distribution for CTD peptides.** Errors are shown as grey area. The associated radii of gyraion (Å) are: 12.86 ± 0.77, 12.86 ± 0.85, 12.77 ± 0.85 for S5S7, 11.82 ± 0.96, 11.72 ± 1.10, 12.02 ± 1.09 for pS5S7, 12.51 ± 0.95, 11.93 ± 0.90, 11.95 ± 0.82 for S5N7, and 12.78 ± 0.45, 12.22 ± 0.42, 12.39 ± 0.41.

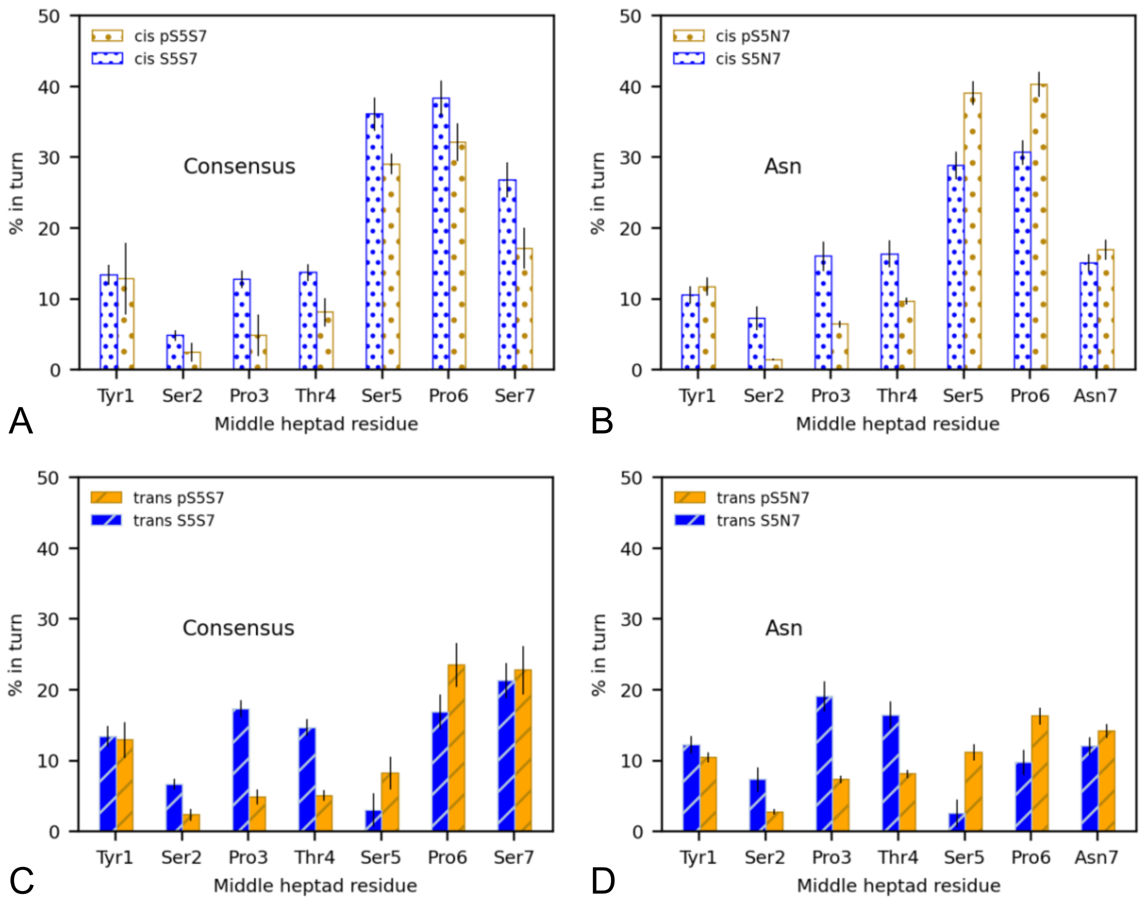


**Figure S6. Percent turn content of middle heptad residues for specific Pro6 isomerization states, related to Figure 8.** (A) and (B) compare the unphosphorylated and phosphorylated ensembles of the consensus and Asn variant respectively, when Pro6 is in the *cis* conformation. (C) and (D) compare the unphosphorylated and phosphorylated ensembles of the consensus and Asn variant respectively, when Pro6 is in the *trans* conformation.

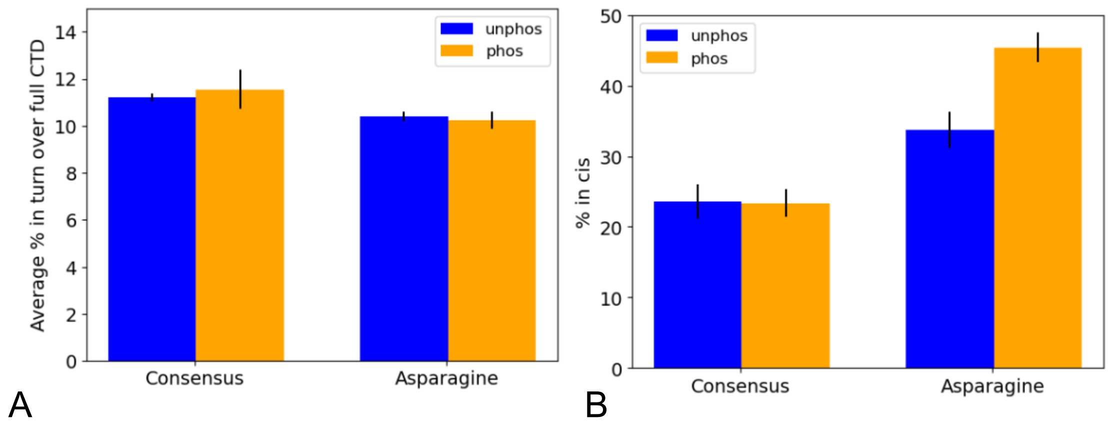


**Figure S7. Composition of turn secondary structure and Pro6 *cis* conformation over entire CTD.** (A) Percent turn secondary structure over all residues of the simulated CTD structure, comparing the unphosphorylated and phosphorylated ensembles for both consensus and Asn variant sequences. (B) Percent Pro6 in the *cis* conformation across all three heptads of the CTD, comparing the unphosphorylated and phosphorylated ensembles for both consensus and Asn variant sequences.

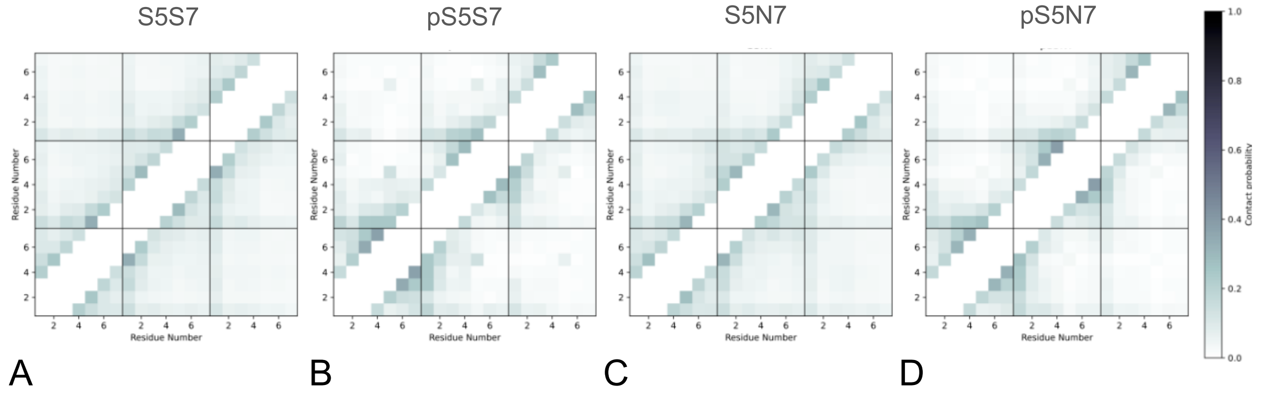


**Figure S8. Contact maps for each CTD construct, related to Figure 10.** (A) Unphosphorylated consensus (B) Phosphorylated consensus (C) Unphosphorylated Asn variant (D) Phosphorylated Asn variant.

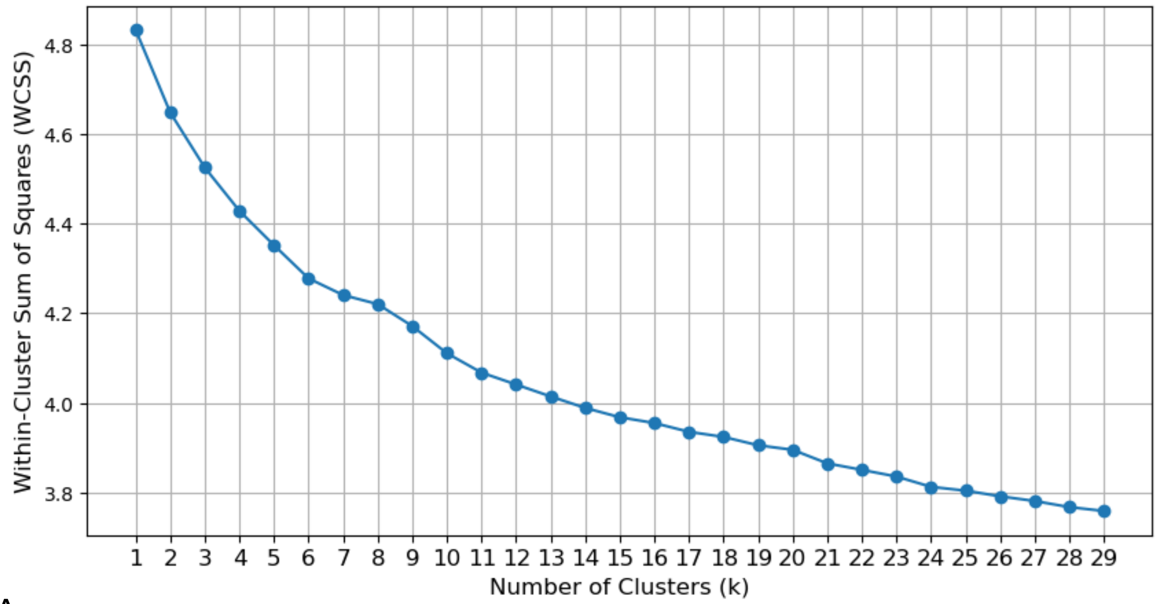


A

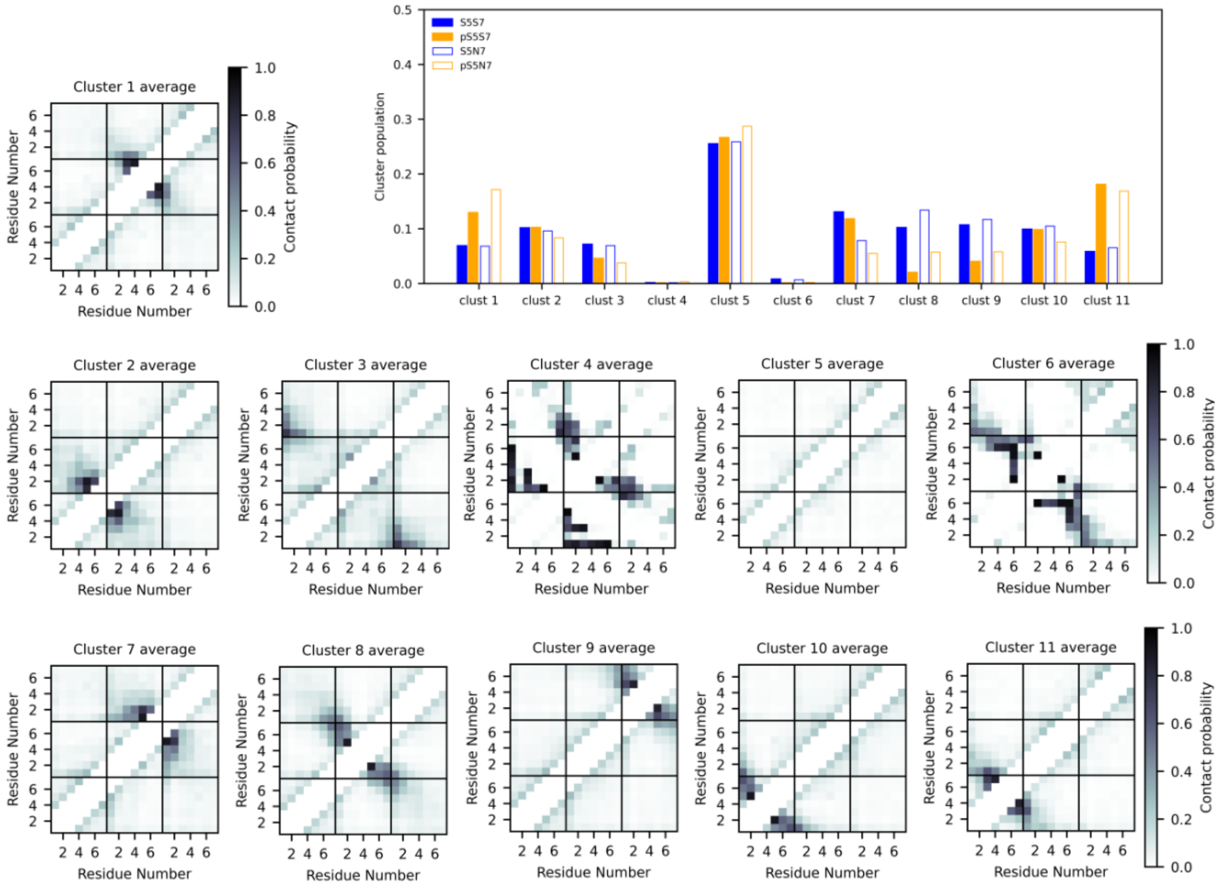


B

**Figure S9. Clustering methodology.** (A) Plot of WCSS versus number of clusters input into *K*-means clustering algorithm. Multiple *k*-values were considered in the "elbow" region, but *k* = 11 clusters revealed several distinct clusters. (B) Average contact maps for each of the 11 clusters. The cluster population plot is displayed again for reference.

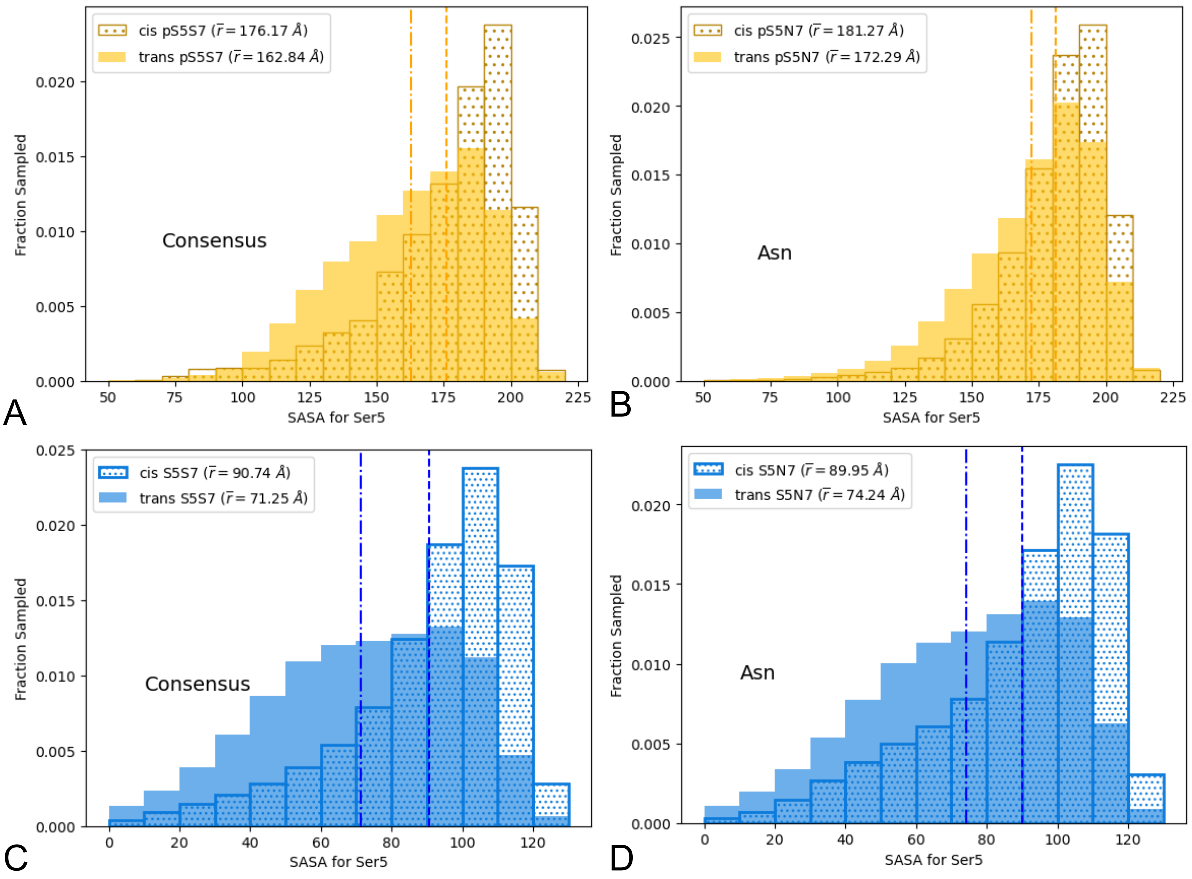


**Figure S10. Solvent-accessible surface area (SASA) histograms.** Dashed lines represent the mean of the *cis*-Pro6 data and dash-dotted lines represent the mean of the *trans*-Pro6 data. (A) Phosphorylated consensus (B) Phosphorylated Asn variant (C) Unphosphorylated consensus (D) Unphosphorylated Asn variant

**Table S1. SAXS analyses on CTD peptides**

| **S5S7** | | | | | | |
|---|---|---|---|---|---|---|
| Replicate | $R_g$ (Å) Guinier | Range (points) | I(0) | $R_g$ (Å) from P(r) | Range (points) | Dmax (Å) |
| 1 | 11.85 ± 0.17 | 16-743 | 0.76 ± 0.0064 | *12.86 +/- 0.77* | 16-743 | *51* |
| 2 | 11.05 ± 0.12 | 20-743 | 0.83 ± 0.0062 | *12.86 +/- 0.85* | 20-743 | *54* |
| 3 | 12.02 ± 0.14 | 26-743 | 0.84 ± 0.0060 | *12.77 +/- 0.85* | 26-743 | *51* |
| **pS5S7** | | | | | | |
| Replicate | $R_g$ (Å) Guinier | Range (points) | I(0) | $R_g$ (Å) from P(r) | Range (points) | Dmax (Å) |
| 1 | 10.88 ± 0.14 | 44-743 | 0.92 ± 0.0073 | *11.82 +/- 0.96* | 44-743 | *41* |
| 2 | 10.75 ± 0.13 | 45.743 | 1.06 ± 0.0079 | *11.72 +/- 1.10* | 45.743 | *40* |
| 3 | 10.95 ± 0.11 | 46-743 | 1.05 ± 0.0069 | *12.02 +/- 1.09* | 46-743 | *43* |
| **S5N7** | | | | | | |
| Replicate | $R_g$ (Å) Guinier | Range (points) | I(0) | $R_g$ (Å) from P(r) | Range (points) | Dmax (Å) |
| 1 | 11.24 ± 0.14 | 27-743 | 0.92 ± 0.0068 | *12.51 +/- 0.95* | 27-743 | *58* |
| 2 | 11.24 ± 0.15 | 27-743 | 0.89 ± 0.0058 | *11.93 +/- 0.90* | 27-743 | *47* |
| 3 | 11.27 ± 0.14 | 21-743 | 0.81 ± 0.0060 | *11.95 +/- 0.82* | 21-743 | *43* |
| **pS5N7** | | | | | | |
| Replicate | $R_g$ (Å) Guinier | Range (points) | I(0) | $R_g$ (Å) from P(r) | Range (points) | Dmax (Å) |
| 1 | 11.18 ± 0.23 | 19-743 | 0.44 ± 0.0053 | *12.78 +/- 0.45* | 19-743 | *48* |
| 2 | 11.16 ± 0.24 | 14-743 | 0.42 ± 0.0053 | *12.22 +/- 0.42* | 14-743 | *44* |
| 3 | 11.48 ± 0.29 | 30-743 | 0.40 ± 0.0062 | *12.39+/- 0.41* | 30-743 | *43* |
